# The human and mouse dendritic cell receptor DCIR binds to LRP1 N-glycans containing terminal galactoses, including the α-Gal antigen

**DOI:** 10.1101/2024.05.20.594848

**Authors:** Benjamin B. A. Raymond, Tamara Sneperger, Stella Rousset, Nelly Gilles, Aurélie Bally, Alexandre Stella, Agnes L. Hipgrave Ederveen, Julien Marcoux, Giulia Trimaglio, Michel Thépaut, Frederic Lagarrigue, Odile Burlet-Schiltz, Manfred Wuhrer, Franck Fieschi, Pascal Demange, Olivier Neyrolles, Yoann Rombouts

## Abstract

The dendritic cell immunoreceptor (DCIR) is a C-type lectin receptor expressed by myeloid cells that plays a key immunoregulatory role in a wide range of diseases, from inflammation to cancer. However, the ligand(s) of DCIR remain(s) unknown, hampering our understanding of the exact function of this immune receptor. Here, we found that both human DCIR and mouse DCIR1 bind specifically to the low-density lipoprotein receptor-related protein 1 (LRP1), a heavily glycosylated receptor mediating the clearance of various molecules from the extracellular matrix and apoptotic cells. This interaction is mediated by galactose-terminated biantennary complex-type N-glycans, including those carrying the immunogenic α-Gal epitope. Our study provides a deeper understanding of the role of DCIR in immune regulation and its potential impact on a range of immune disorders, highlighting its specificity in ligand recognition which is crucial for developing therapeutic strategies.

## INTRODUCTION

Myeloid cells constantly monitor the tissues in which they reside to detect and respond quickly to foreign bodies or any internal damage. For that purpose, they express a plethora of pattern recognition receptors (PRRs), including C-type lectin receptors (CLRs).^1, 2^ CLRs were originally named for their ability to bind carbohydrates in a Ca^2+^-dependent manner via their C-type lectin domain (CTLD). In this regard, many CLRs bind carbohydrates covering microbes and play an essential role in the defense against infections. However, it is now clear that some CLRs also interact with non-carbohydrate structures such as proteins released by damaged cells, and regulate immunity in a broad spectrum of diseases ranging from inflammatory and autoimmune disorders to cancer.^1, 2^

Most CLRs carry, or are coupled to, an intracellular immunoreceptor tyrosine-based activation motif (ITAM) whose downstream signaling pathways usually lead to the production of pro-inflammatory mediators by myeloid cells. Few CLRs possess an immunoreceptor tyrosine inhibitor motif (ITIM) that, when phosphorylated, associates with phosphatases 1 and 2 containing the Src region homology domain (SHP1 and SHP2), acting as an intrinsic brake on various immune cell signaling pathways.

The dendritic cell immunoreceptor (DCIR) is an ITIM-bearing CLR belonging to the Dectin-2 family.^3^ Humans possess a single gene encoding DCIR, *CLEC4A,* expressed in several subtypes of myeloid cells, including conventional and plasmacytoid dendritic cells (DCs), macrophages, monocytes, and granulocytes, and B cells. Mice express four homologues of hDCIR, encoded by the *Clec4a1-4* genes, yet only mDCIR1 (CLEC4A2) and mDCIR2 (CLEC4A4) contain an ITIM motif. mDCIR1 is often considered the functional homologue of hDCIR due to its greater sequence similarity and comparable expression profile on myeloid cells, unlike mDCIR2 whose expression is restricted to CD8α negative conventional DCs.^4^ The study of mDCIR1-KO (*Clec4a2^-/-^*) C57BL/6 mice has highlighted the important immunoregulatory role of this CLR in a plethora of immune disorders, including chronic inflammatory diseases, autoimmunity, allergy, cancer and infectious diseases.^5–22^ A recurrent phenotype observed in mDCIR1-KO mice is the increase of the adaptive immune response, providing better immunity against microbes but leading to hyperinflammation or the development of autoimmune diseases.^8–11, 15^ The increased adaptive immune response in *mDCIR1*-KO mice was explained notably by dysfunctions in DCs in these mice, including their hyperproliferation in response to GM-CSF, an attenuated response to type I interferons (IFN) counterbalanced by an overproduction of IL-12p70, a key cytokine for the induction of T cell response, and an overall enhanced antigen-presentation ability.^7–11, 15^ *In vitro* studies have also established that hDCIR and mDCIR1 on DCs and macrophages may counteract the pro-inflammatory response following stimulation of Toll-like receptors (TLRs), especially endosomal TLR7/8/9.^6, 8, 20, 23–26^ Contrary to these findings, several studies have also found a reduced infiltration of innate immune cells and cytokine/chemokine production in mDCIR1-KO mice during acute inflammation or infection, although the underlying mechanisms remain unclear.^5–7, 13, 19, 22^

One of the major difficulties that hinders our understanding of the function of DCIR is that its ligand(s) have still not been clearly identified. hDCIR has been repeatedly shown to recognize certain glycans in a Ca^2+^-dependent manner, but the glycosylation motifs recognized by this CLR vary between studies. hDCIR has been shown to interact with fucose- or mannose-containing carbohydrates as well as N-linked glycans containing terminal N-acetylglucosamine (GlcNAc), galactose (Gal) or sialic acid residues.^9, 16, 27–31^ Notably, all the sugar motifs identified as hDCIR ligands are carried by proteins in the human body, supporting the idea that this CLR recognizes endogenous glycoproteins that have yet to be characterized. In this respect, hDCIR contains an N-glycosylation site and could interact with itself, as described in cells overexpressing hDCIR.^25, 28, 29^ In the absence of known glycoprotein ligand(s) for hDCIR, it remains unclear whether mDCIR1 recognizes similar ligands and therefore whether the mouse model is well suited to studying the role of this CLR in immunity.

Here, we identified the low-density lipoprotein receptor-related protein 1 (LRP1), a highly glycosylated receptor, as a main endogenous ligand of mDCIR1 and hDCIR. We further showed that LRP1 recognition by DCIR depends on LRP1 N-glycans, characterized the N-glycosylation profiles of mouse and human LRP1, and showed that both mDCIR1 and hDCIR bind specifically to galactose-terminated biantennary complex-type N-glycans including those carrying the highly immunogenic α-gal epitope. Finally, we obtained the high-resolution crystal structure of hDCIR bound to the biantennary N-glycan carrying terminal α-gal epitopes and, by means of mutagenesis experiments, we were able to define the structural basis of hDCIR binding to its ligand.

## RESULTS

### Mouse myeloid cells express both mouse DCIR1 and its ligand(s)

In an attempt to identify cells that express mDCIR1 ligand(s), we produced a stable recombinant extracellular domain (ECD) of mDCIR1 and also generated a mDCIR1-ECD mutated in the calcium-binding site (ΔECD; E197A) to be used as a negative control. Both the mDCIR1-ECD and mDCIR1-ΔECD were biotinylated, coupled to fluorescent streptavidin and used in flow cytometry experiments to reveal cells expressing mDCIR1 ligand(s) in lymphoid organs (*i.e.*, spleen and thymus), non-lymphoid organs (*i.e.*, lungs and bone marrow) and in blood of mice (Figure 1A; Gating strategy in Figure S1). We observed that mDCIR1-ECD binds preferentially to myeloid cells, namely neutrophils, macrophages, DCs and monocytes, but weakly to lymphocytes in all tissues analyzed (Figures 1B-F). Since myeloid cells express mDCIR1 (Figure S2) and since it has been previously proposed that mDCIR1 may bind to itself, we performed the same experiment on tissues from mDCIR1-deficient mice. We found that mDCIR1-ECD binds to mDCIR1-KO cells in a similar manner to WT cells, which does not support the hypothesis that mDCIR1 interacts primarily with itself (Figures 1B-F). We concluded that one or more endogenous ligands of mDCIR1 are expressed at the cell surface of macrophages, DCs, monocytes and neutrophils, which can be used for further biochemical characterization.

**Figure 1:**
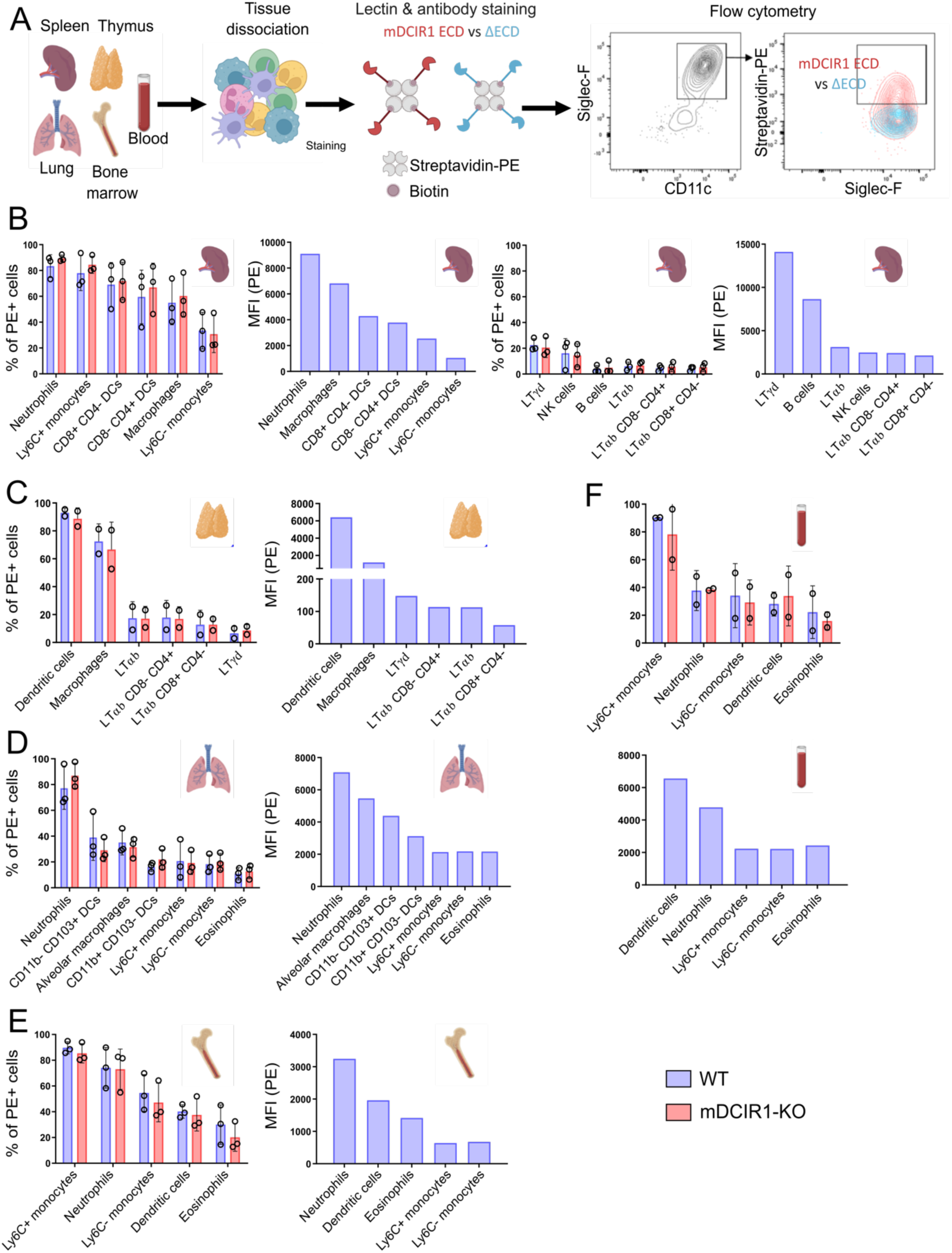
Flow cytometry analysis identifies myeloid innate immune cells as carriers of mDCIR1 ligand(s). (A) Schematic of the strategy used for flow cytometric analysis of cells expressing mDCIR1 ligand(s) in different tissues of WT and mDCIR1-KO mice (Created with BioRender.com). (B) Percentage (left panels) of cells stained with biotinylated mDCIR1-ECD coupled to streptavidin--Phycoerythrin (Streptavidin-PE) cells and median fluorescent intensity (MFI; right panels) in the spleen of WT and mDCIR1-KO mice. Only the MFI of WT cells is shown. Gating strategies used for the identification of cell subtypes are depicted in figure S1. (C) Same as (B) but for the thymus. (D) Same as (B) but for the lung. (E) Same as (B) but for the bone marrow. (F) Same as (B) but for blood.

### Mouse DCIR1 binds to LRP1 α-chain in macrophages and dendritic cells

Because mDCIR1 has previously been shown to impact macrophage and DC functions and since these cells express mDCIR1 ligand(s) in all of the organs analyzed, we set out to identify the mDCIR1 ligand(s) in these cell types. We first confirmed that mDCIR1-ECD, but not mDCIR1-ΔECD, binds to bone marrow-derived macrophages (BMDM) and dendritic cells (BMDC) (Figure 2A). We also demonstrated that the binding of mDCIR1-ECD to these cells requires divalent ions, most probably Ca^2+^, as it can be inhibited by the addition of EDTA and EGTA chelating agents (Figure 2A). In order to determine whether mDCIR1 binds to one or several proteins on BMDM and BMDC, we lysed these cells, resolved their proteins on SDS-PAGE and performed lectin blots using the biotinylated mDCIR1-ECD, conjugated to streptavidin-HRP. While we observed no binding of mDCIR1-ECD within the range of 10-250 kDa, we observed a strong band of reactivity well above 250 kDa in both cell types (Figure 2B). Importantly, no binding was evidenced with mDCIR1-ΔECD (Figure S3A). Next, we attempted to isolate the ligand(s) of mDCIR1 from BMDM through mDCIR1-ECD affinity pulldown experiments. The bound material was subsequently eluted via an excess of Ca^2+^ chelator (*i.e*., EGTA) and analyzed by proteomics (Figure 2C). As negative controls, similar experiments were performed either using mDCIR1-ΔECD instead of mDCIR1-ECD (Figure 2C), or in the presence of EGTA throughout the experiment to inhibit ligand recognition by the lectin (Figure S3B). Mass spectrometry analysis of pull-down proteins revealed a strong enrichment of the low-density lipoprotein receptor-related protein 1 (LRP1) and its binding partners, namely the LDL Receptor Related Protein Associated Protein 1 (LRPAP1) and Thrombospondin 1 (THBS1) in the mDCIR1-ECD eluate as compared to the control eluates (Figures 2C and S3B; Tables S1). LRP1 is a receptor composed of an extracellular α-chain of 515 kDa and a transmembrane β-chain of 85 kDa. In a duplicate affinity pull-down experiment, we confirmed the enrichment of LRP1 α- and β-chains with mDCIR1-ECD but not with mDCIR1-ΔECD by western blotting (Figure S3C). These experiments did not reveal enrichment of LRPAP1, and led to only slight enrichment of THBS1 (Figure S3C). Based upon proteins with the greatest number of unique peptides identified by proteomics and the high molecular weight bands observed by lectin blot, we hypothesized that mDCIR1 binds to LRP1 α-chain. By immunoprecipitating LRP1 and then performing lectin blot analysis, we confirmed strong binding of mDCIR1-ECD, but not mDCIR1-ΔECD, to the LRP1 α-chain while no binding was observed to the LRP1 β-chain (Figure 2D). The identity of the LRP1 α-chain was further assessed by immunoblotting and tandem mass spectrometry (Figure 2D & Tables S2). We next generated *Lrp1*-KO macrophages using CRISPR/Cas9 editing in ER-Hoxb8 conditionally immortalized hematopoietic progenitors to analyze the binding of mDCIR1-ECD by flow cytometry (Figures 2E and S4). Compared with control cells, we observed a clear decrease in mDCIR1-ECD binding to *Lrp1*-KO cells. Altogether, these data demonstrate that LRP1 is the main ligand of mDCIR1 in macrophages and DCs.

**Figure 2:**
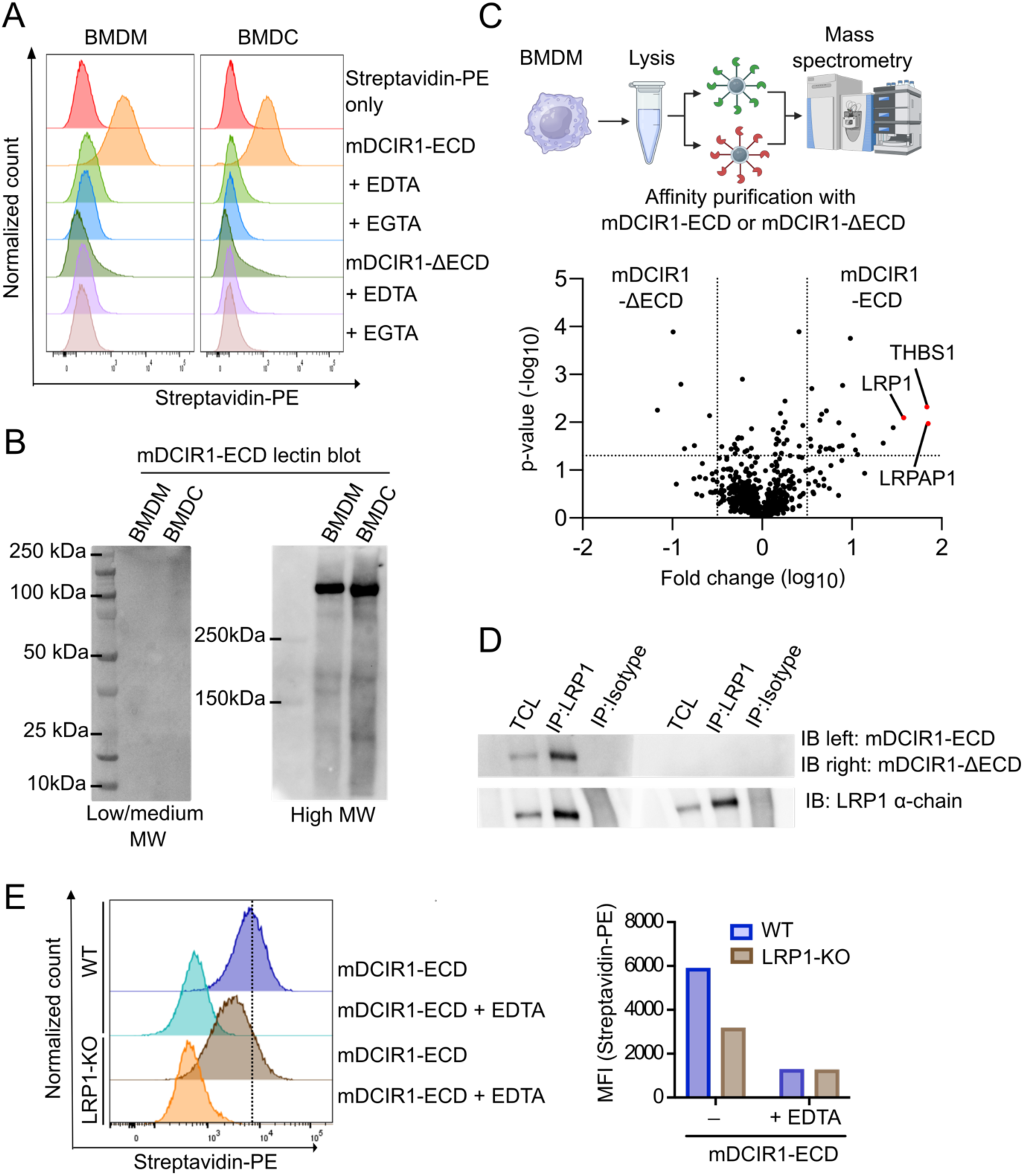
LRP1 is a main glycoprotein ligand of mDCIR1. (A) Flow cytometry histograms of the binding of biotinylated mDCIR1-ECD and mDCIR1-βECD coupled to Streptavidin-Phycoerythrin (Streptavidin-PE) to bone marrow-derived macrophages (BMDM) and dendritic cells (BMDC), in the absence or presence of EDTA and EGTA chelating agents. (B) Lectin blot examination of mDCIR1 ligand(s) extracted from BMDM and BMDC using biotinylated mDCIR1-ECD coupled to streptavidin-HRP. (C) Schematic (Top) of the enrichment of mDCIR1 ligand(s) from BMDM by affinity purification using biotinylated mDCIR1-ECD (or βECD as negative control) coupled to streptavidin beads and their subsequent identification by proteomics (Created with BioRender.com). Volcano Plot (bottom) from quantitative mass spectrometry analyses of proteins differentially enriched (X-axis in log_10_) by affinity purification with mDCIR1-ECD versus mDCIR1-βECD as a function of statistical significance (Y-axis in -log10). Dashed line marks the threshold limit (P-value = 0.5; Fold change = +/- 5). (D) Analysis of LRP1 immunoprecipitated from BMDM by western blot and lectin blot using an anti-LRP1 alpha chain antibody and biotinylated mDCIR1-ECD or mDCIR1-βECD (coupled to streptavidin-HRP) respectively. (E) Flow cytometry histograms (left panel) and median fluorescent intensity (MFI; right panel) of the binding of biotinylated mDCIR1-ECD (coupled to Streptavidin-PE) to WT and LRP1-deficicent HoxB8-derived macrophages, in the absence or presence of EDTA.

### Human DCIR binds to LRP1 α-chain in macrophages and dendritic cells

Mouse DCIR1 is typically studied as the homolog of hDCIR; however, it is not known whether these two CLRs recognize the same ligand(s). We therefore investigated whether hDCIR could also bind to LRP1 on human macrophages and DCs. To this end, we produced the extracellular domain of human DCIR (hDCIR-ECD) and evaluated its capacity to bind to human blood monocyte derived-macrophages (mo-Mac) and DCs (mo-DC) by flow cytometry. We observed that hDCIR-ECD binds to mo-Mac and mo-DC in a Ca^2+^-dependent manner (Figure 3A). We then carried out affinity pulldown experiments using hDCIR-ECD to capture potential ligand(s) on mo-Mac and mo-DC followed by proteomics analysis (Figure 3B). As a negative control, pulldown experiments were performed in the presence of EGTA during all steps of the experiment. In both cell types, LRP1 was highly enriched by the hDCIR-ECD affinity pulldown (Figure 3B and Table S1). Furthermore, immunoprecipitation of LRP1 from mo-Mac and mo-DC followed by lectin blot confirmed the direct binding of hDCIR to LRP1 α-chain (Figure 3C). Gene silencing of LRP1 in mo-Mac using siRNAs abolished hDCIR-ECD binding to cells (Figure S5A). However, a similar reduction was observed when cells were treated with non-targeted control siRNAs. This may be explained by the fact that siRNA transfection by itself strongly reduces LRP1 expression as quantified by RT-qPCR (Figure S5B). To circumvent this issue, we analyzed hDCIR-ECD binding to WT and *Lrp1*-KO murine macrophages and found that LRP1 deficiency completely abolished lectin binding (Figure 3D). Altogether, these results demonstrate that LRP1 α-chain is also a ligand for hDCIR.

**Figure 3:**
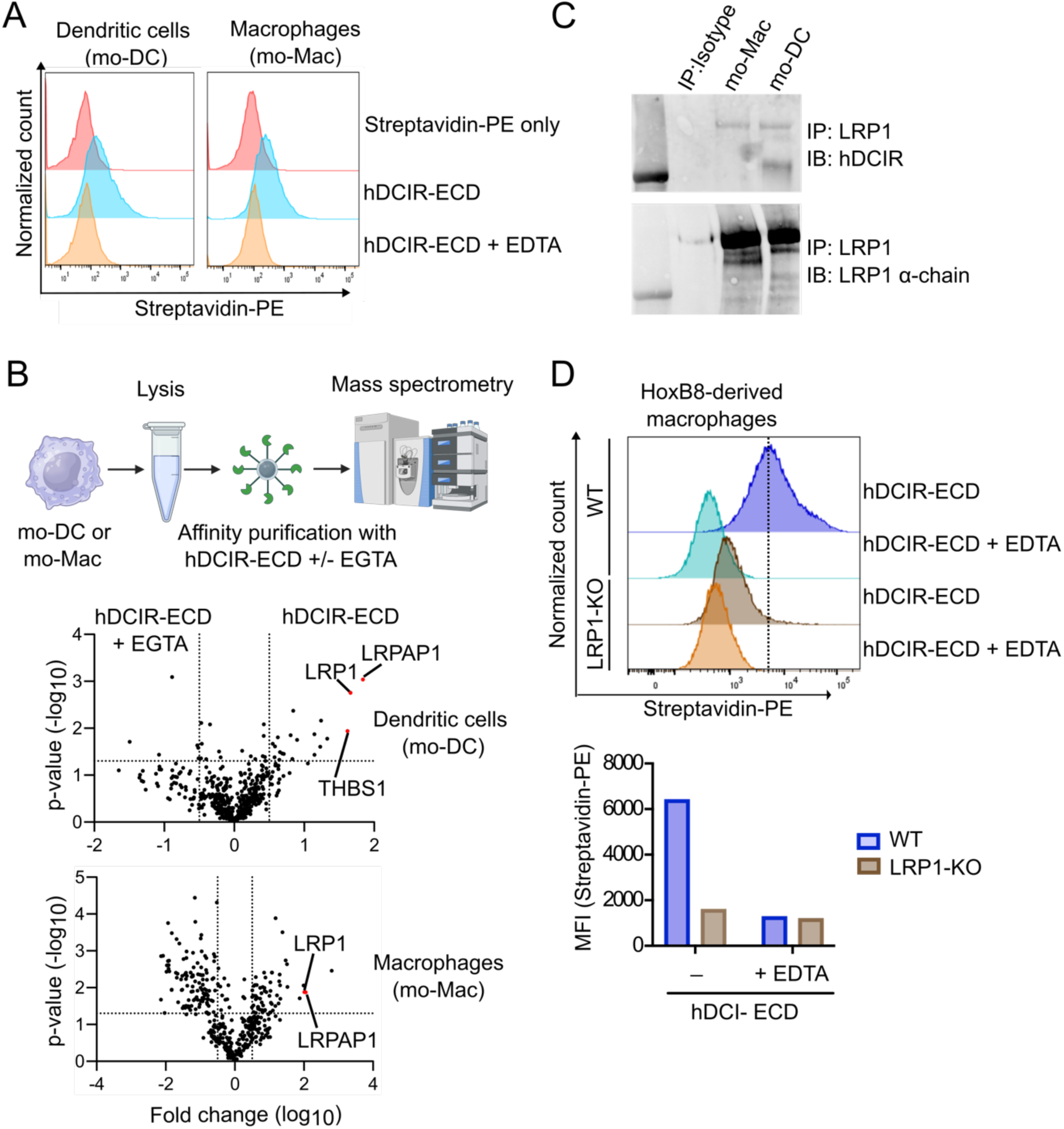
LRP1 is also a major ligand of hDCIR. (A) Flow cytometry histograms of the binding of biotinylated hDCIR-ECD coupled to Streptavidin-PE to human blood monocyte-derived dendritic cells or macrophages (mo-DC or mo-Mac), in the absence or presence of EDTA chelating agent. (B) Schematic (Top) of the enrichment of hDCIR ligand(s) from mo-DC or mo-Mac by affinity purification using biotinylated hDCIR-ECD (or hDCIR-ECD in the presence of EGTA as negative control) coupled to streptavidin beads and their subsequent identification by proteomics (Created with BioRender.com). Volcano Plots (middle and bottom) from quantitative mass spectrometry analyses of proteins differentially enriched (X-axis in log_10_) by affinity purification with hDCIR-ECD versus hDCIR-ECD plus EGTA as a function of statistical significance (Y-axis in -log10). Dashed line marks the threshold limit (P-value = 0.5; Fold change = +/- 5). (C) Examination of LRP1 immunoprecipitated from mo-Mac and mo-DC by western blot and lectin blot using an anti-LRP1 alpha chain antibody and hDCIR-ECD. (D) Flow cytometry histograms (upper panel) and median fluorescent intensity (MFI; bottom panel) of the binding of biotinylated hDCIR-ECD (coupled to Streptavidin-PE) to WT and LRP1-deficicent HoxB8-derived macrophages, in the absence or presence of EDTA.

### Both mDCIR1 and hDCIR bind to biantennary complex-type N-glycans with terminal galactose residues

Given that the interaction of mDCIR1 and hDCIR to myeloid cells requires the presence of Ca^2+^, we hypothesized that mDCIR1 and hDCIR may bind to LRP1 α-chain in a glycan-dependent manner. LRP1 is a highly glycosylated protein with over 45 putative N-linked glycosylation sites, of which more than twenty have been confirmed, and several O-linked glycans.^32–34^ As DCIR has previously been shown to bind mostly to N-glycan structures, we first removed the N-glycans from immunoprecipitated LRP1, using Peptide:N-glycosidase F (PNGase F), and evaluated by lectin blot whether the binding of mDCIR1-ECD was affected (Figure 4A). As expected, the removal of N-glycans strongly decreased the molecular weight of LRP1 α-chain. Importantly, it also completely abolished binding of mDCIR1-ECD to LRP1 α-chain (Figure 4A). With the aim to characterize the N-glycan epitope recognized by mDCIR1 on LRP1, we purified LRP1 α-chains from BMDM and human serum, and analyzed their N-glycan structures by capillary electrophoresis-electrospray ionization-mass spectrometry (CE-ESI-MS) (Figure S6A and Table S3).^35^ Both mouse and human LRP1 carried almost exclusively biantennary complex-type N-glycans with either terminal galactose or sialic acid (N-acetylneuraminic acid) residues (Figure 4B). The latter were mainly α2,6-linked sialic acids as determined by linkage-specific derivatization of sialic acids.^35^ Of note, tandem mass spectrometry (MS/MS) and antibody staining showed that mouse LRP1 N-glycans exhibit the terminal Galα1-3Galβ1-4GlcNAc (α-Gal) epitope (Figures 4B and S6), a unique carbohydrate antigen abundantly expressed on glycoconjugates of most mammals yet absent in humans.^36, 37^ To determine which of these N-glycan structures are ligand(s) for hDCIR and mDCIR1, we performed an enzyme-linked lectin assay using neo-glycoproteins generated by covalent coupling of biantennary complex-type N-glycans with either terminal GlcNAc, Gal, α-Gal or α2,6-linked sialic residues to bovine serum albumin (BSA). We found that both hDCIR-ECD and mDCIR1-ECD strongly interact with biantennary complex-type N-glycans with terminal galactose residues, including those carrying the α-Gal epitope, but poorly or not at all with agalactosylated or sialylated N-glycans (Figure 4C). As expected, lectin binding to these glycan structures was Ca^2+^-dependent, being completely abolished in the presence of EDTA. Further analysis by flow cytometry confirmed that hDCIR-ECD and mDCIR1-ECD bound to N-glycans harboring terminal galactoses as expressed in the Chinese Hamster Ovary (CHO) glycosylation mutant Lec2 cells, which are defective in Golgi transport of CMP-sialic acid, but not in Lec8 cells, which lack a functional UDP-Galactose transporter (Figure 4D).

**Figure 4:**
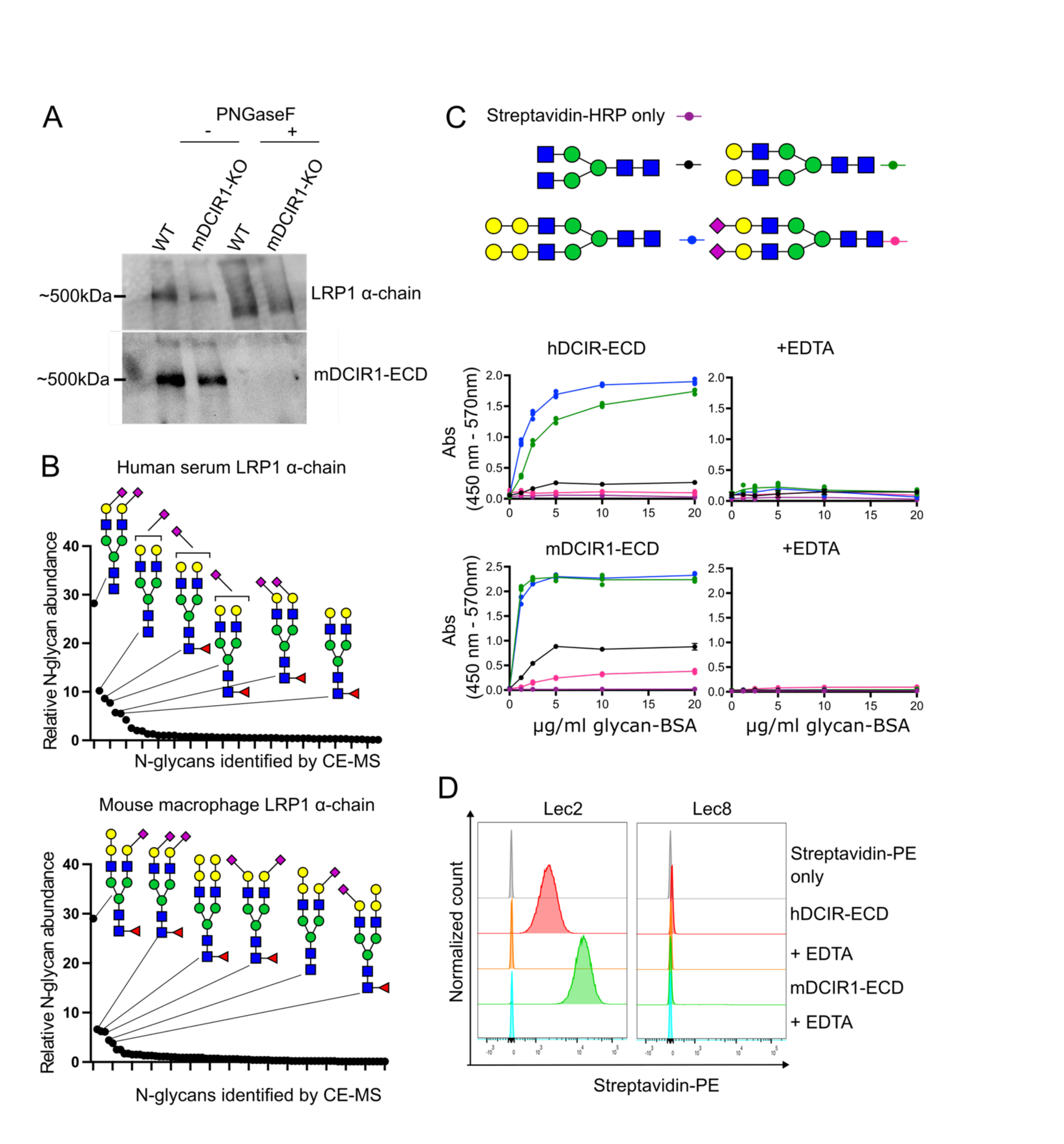
mDCIR1 and hDCIR interact with galactose-terminated biantennary biantennary N-glycans, as carried by LRP1. (A) Western blot and lectin blot analysis of LRP1 immunoprecipitated from BMDM, before or after PNGase F-catalyzed N-glycan removal, using an anti-LRP1 α-chain antibody and biotinylated mDCIR1-ECD coupled to streptavidin-HRP. (B) Most abundant N-glycan carried by LRP1 from human serum and mouse BMDM as identified by CE-ESI-MS analysis. Green circle: mannose, yellow circle: galactose, blue square: N-acetylglucosamine, right pointing purple diamond: α2,6-linked N-acetylneuraminic acid/sialic acid, left pointing purple diamond: α2,3-linked N-acetylneuraminic acid/sialic acid and red triangle: fucose. (C) Binding of hDCIR-ECD and mDCIR1-ECD (coupled to streptavidin-HRP) to different biantennary complex-type N-glycans chemically linked to bovine serum albumin (BSA), in the absence or presence of EDTA, as measured by lectin ELISA. Green circle: mannose, yellow circle: galactose, blue square: N-acetylglucosamine, purple diamond: α2,6-linked N-acetylneuraminic acid/sialic acid. (D) Flow cytometry histograms of the binding of hDCIR-ECD and mDCIR1-ECD (coupled to streptavidin-PE), in the absence or presence of EDTA, to Lec2 or Lec8 cell lines harboring biantennary complex-type N-glycans containing terminal galactose residues, or terminal N-acetylglucosamine respectively

### Structural basis of hDCIR interaction with terminally galactosylated biantennary complex-type N-glycans

Since the discovery of hDCIR in the late 1990s, only two medium-resolution (∼ 3.0 Å) X-ray crystallography structures of the hDCIR C-type lectin domain (CTLD) were obtained.^30^ One of these structures corresponds to hDCIR CTLD interacting in a Ca^2+^-dependent manner with an oligosaccharide mimicking a biantennary N-glycan unit (Galβ1-4GlcNAcβ1-2Manα1-3[GlcNAcβ1-2Manα1-6]Manα1-O-tert-butyldimethylsilyl), one antenna of which contains a terminal galactose residue while the other is non-galactosylated. Based on this structure, it was concluded that hDCIR interacts with mannose residues in its Ca^2+^-binding site and that GlcNAcβ1-2Man, with or without terminal galactose, on both antennae is the main epitope recognized by hDCIR. On the contrary, our data show that the binding of hDCIR to biantennary N-glycans requires the presence of terminal galactoses. Human DCIR contains a carbohydrate recognition motif composed of Glu-Pro-Ser (EPS) similar to the prototypical Glu-Pro-Asn (EPN) motif, which is generally considered to display mannose specificity. However, there are examples of C-type lectins containing either an EPN or an EPS motif that exhibit preferential binding to galactose residues.^38, 39^ To investigate the structural basis of hDCIR binding to its ligands, we crystallized hDCIR-ECD alone or in the presence of biantennary complex-type N-glycans with either terminal galactoses or terminal α-Gal epitopes. Although we obtained crystals in all these conditions, we only collected diffraction-quality X-ray crystallographic data for hDCIR-ECD bound to biantennary N-glycans with terminal α-gal epitopes at a resolution of ∼ 1.9 Å (Figure 5A). The structure of hDCIR-ECD presents a typical C-type lectin fold and was comparable to previously obtained structures of hDCIR CTLD (Figure S7A). We observed a well-defined electron density of antennary Galα1-3Galβ1-4GlcNAcβ1-2Manα1-3 in the primary sugar-binding site of hDCIR, but partial electron density for the rest of the oligosaccharide (Figure 5). We therefore only assigned the density map for Galα1-3Galβ1-4GlcNAcβ1-2Manα in complex with the lectin (Figure 5). The crystal packing of the ligand-lectin complex showed that the two antennae of a given N-glycan were embedded in the sugar-binding site of two different hDCIR units (Figure 5A). A closer look at the hDCIR binding site, notably using a two-dimensional interaction map, highlighted all the interactions between the carbohydrate and hDCIR in the crystal structure of the complex (Figures 5B-C). The calcium ion was chelated by the Glu195 and Ser197 residues of the EPS motif, by the Asn218 and Asp219 residues of the WND motif, as well as by the Glu201. The carbohydrate also participated in the chelation of the calcium ion via the two equatorial hydroxyl groups (3-OH and 4-OH) of the mannose residue. The participation of mannose in the chelation of the calcium ion depends on the orientation of the adjacent GlcNAc and its interaction with the Arg209 and Asn218. hDCIR interacts with the two terminal galactose residues via hydrogen bonds formed between the side chain of Gln226 and the 2-OH and 6-OH groups of the first and terminal galactose residues, respectively. The terminal galactose was also stabilized via a hydrogen bond involving its 4-OH group and the hydroxyl group of Ser124. The two galactoses also interacted with hDCIR via van der Waals interactions involving amino acids Glu160, Ser161 and Arg227. The key amino acids involved in hDCIR binding to Galα1-4Galβ1-4GlcNAcβ1-2Manα1-3 are well conserved in mDCIR1 (*i.e.*, EPS and WND motifs forming the core of the binding site as well as Arg209, Gln226 and Ser124 involved in the interaction with the GlcNAc and galactoses) (Figure S7B). In contrast, other murine homologues of DCIR show variations in their amino acid sequences (Figure S7B). Indeed, mDCIR2 has a prototypical EPN motif, lacks the Ser124 and carries two histidines instead of Arg209 and Glu226. These histidine residues are also present in mDCIR4, which also lacks the Asn218 and Asp219 of the WND motif involved in calcium ion coordination. Finally, mDCIR3 is the most similar to hDCIR and mDCIR1, although it lacks the Asn218 of the WND motif and the Ser124.

**Figure 5:**
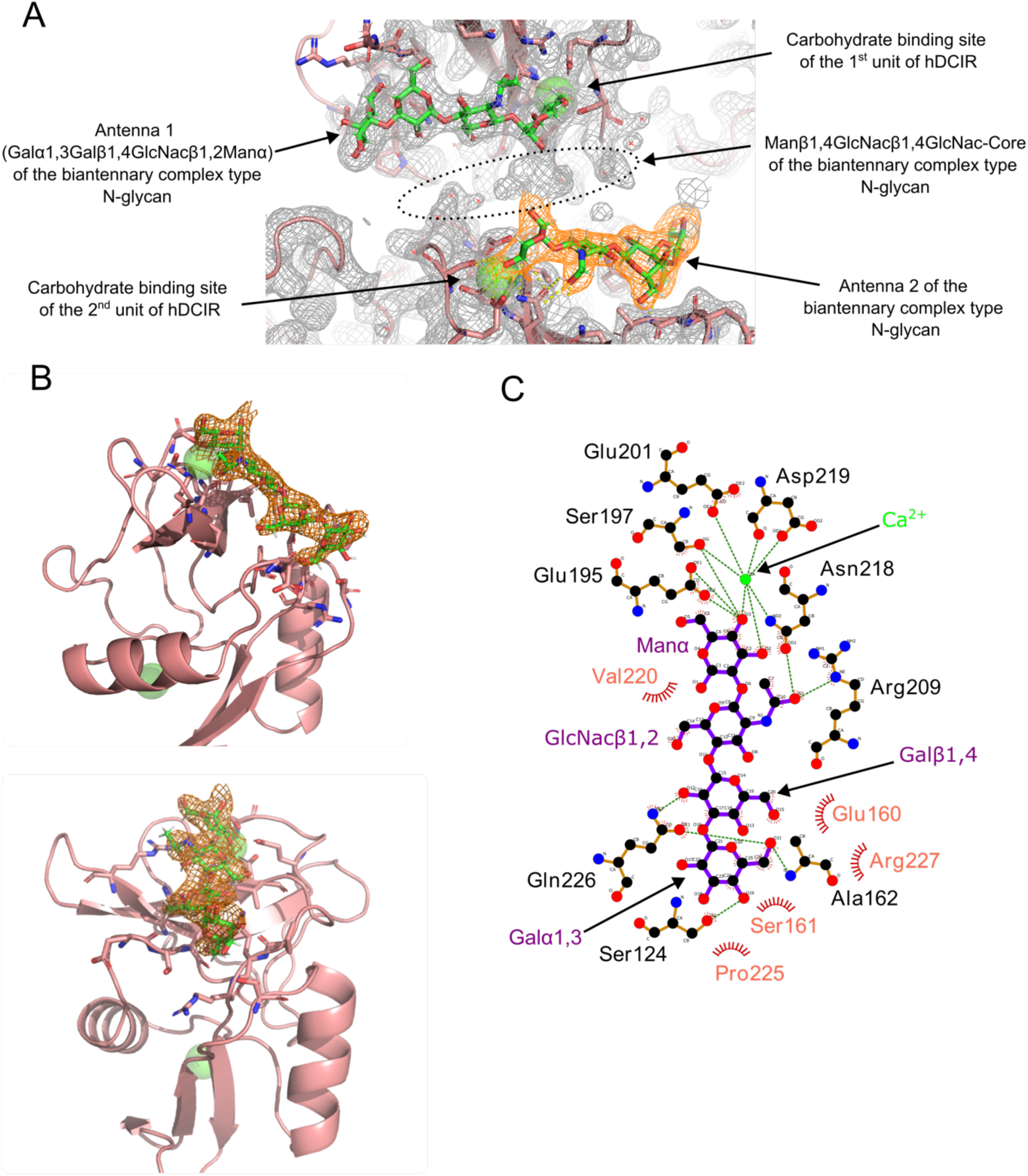
X-ray crystallography revealed the structural basis of hDCIR binding to biantennary complex-type N-glycans containing terminal galactoses. (A) Close-up view of the crystal packing and electron density of the sugar binding sites of two units of hDCIR-ECD interacting with the Galα1-3Galβ1-4GlcNAcβ1-2Manα antennae of a biantennary N-glycan carrying terminal α-Gal epitopes. Protein, carbohydrate, and calcium ions are shown in ribbon (pink), stick (green/red), and sphere (light green) models, respectively. Amino acid residues which interact with the tetrasaccharide and calcium ion are shown by stick models and are labeled. The electron density is represented by the grey or orange mesh. (B) Close-up views of hDCIR-ECD in complex with the Galα1-3Galβ1-4GlcNAcβ1-2Manα antenna of the N-glycan. Protein, carbohydrate, and calcium ions are represented as in (A). The electron density of the tetrasaccharide is represented by the orange mesh. (C) Two-dimensional interaction map of the Galα1-3Galβ1-4GlcNAcβ1-2Manα antenna of biantennary complex type N-glycan bound to hDCIR. The hydrogen bonds and coordination involved in the interaction between the ligand, the calcium ion, and atoms of the protein residues are represented by dotted green lines. Van der Waals contact between the ligand and the protein are represented by notched semicircles.

Based on these data, we hypothesized that terminal galactoses are required for hDCIR binding to biantennary N-glycans due to i) the interaction between the side chain of Gln226 and hydroxyl groups of galactoses and ii) the Ser197 of the EPS motif, which we suspect has a weaker commitment in mannose binding and/or calcium coordination than a Gln in a classical mannose-type EPN motif. To confirm this hypothesis, we mutated the residues S197A (Ser197Asn), Q226A (Gln226Ala) and as control E195A (Glu195Ala; *i.e.*, to abolish the binding to Ca^2+^) of hDCIR-ECD and measured the binding of the lectin mutants to the different glycan structures by enzyme-linked lectin assay. We found that E195A and Q226A mutations completely abolished the binding of hDCIR-ECD to galactose-capped biantennary N-glycans. In contrast, the S197N mutation not only enhanced the binding of hDCIR-ECD to galactosylated biantennary N-glycans, but also conferred binding to non-galactosylated biantennary N-glycans (Figure 6A). Similarly, the E195A and Q226A mutations suppressed hDCIR-ECD binding to Lec2 cells, while the S197N mutation promoted its binding to both Lec2 and Lec8 cells (Figure 6B). Altogether, our data revealed the structural mechanism by which hDCIR binds to galactose-terminated biantennary complex-type N-glycans.

**Figure 6:**
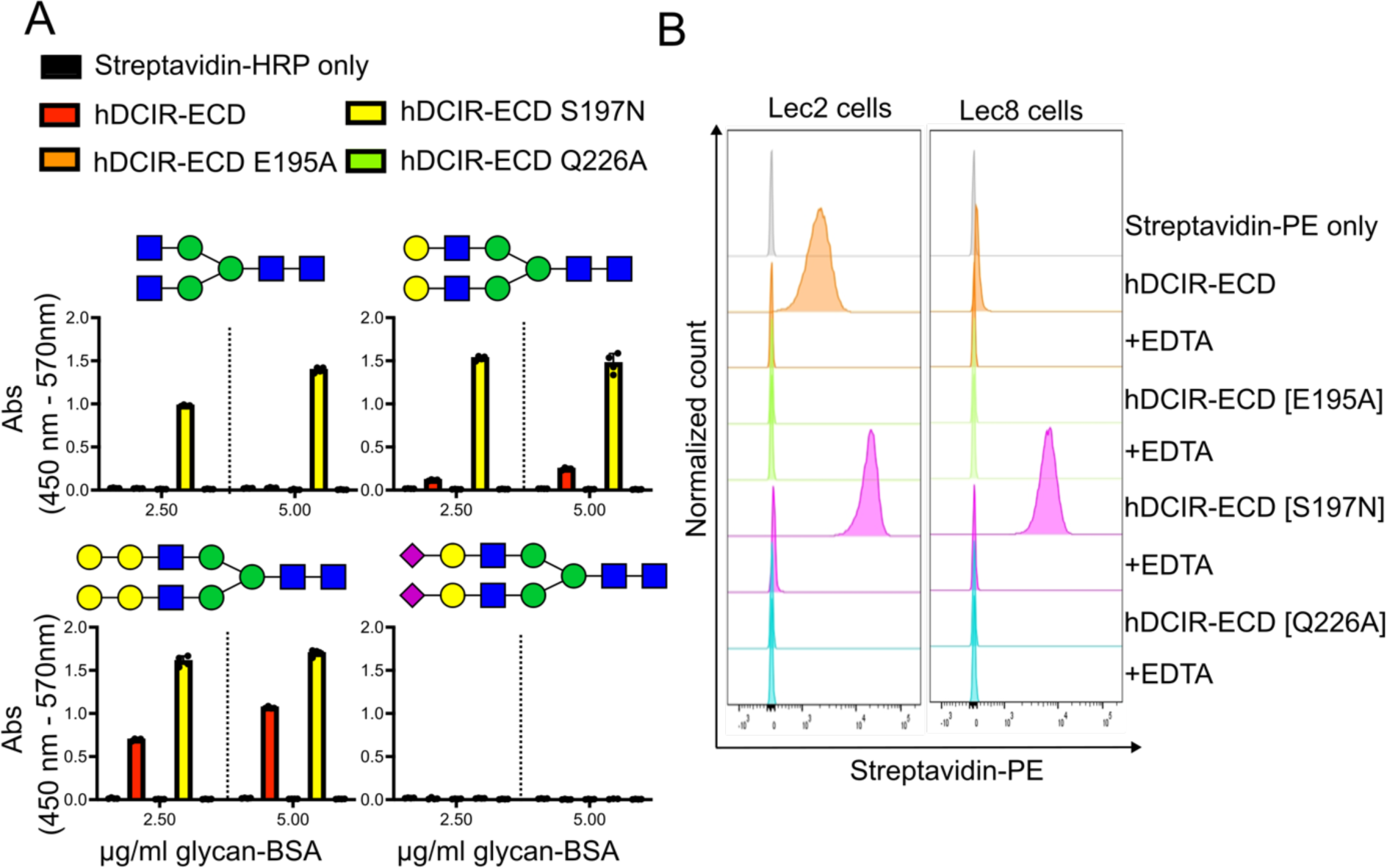
Mutagenesis confirmation of the mechanism for binding of galactose-terminated biantennary N-glycans by hDCIR. (A) Binding of hDCIR-ECD and mutants (coupled to streptavidin-HRP) to different biantennary complex-type N-glycans attached to bovine serum albumin (BSA) as measured by lectin ELISA. Green circle: mannose, yellow circle: galactose, blue square: N-acetylglucosamine, purple diamond: α2,6-linked N-acetylneuraminic acid/sialic acid. (B) Flow cytometry histograms of the binding of hDCIR-ECD and mutants (coupled to streptavidin-PE), in the absence or presence of EDTA, to Lec2 or Lec8 cell lines. Green circle: mannose, yellow circle: galactose, blue square: N-acetylglucosamine, violet diamond: N-acetylneuraminic acid/sialic acid. Linkages between monosaccharides are indicated.

## DISCUSSION

Although literature data suggest that hDCIR interacts with one or more endogenous glycoprotein ligands, these remained unidentified to date. Consequently, it was impossible to determine whether mDCIR1 (Clec4a2), the closest murine homologue of hDCIR, recognized the same ligand and whether mDCIR1-deficient mice constituted a relevant model for characterizing the immune function of hDCIR. Here, we identified the highly glycosylated cell-surface protein LRP1, expressed notably by macrophages and DCs, as a main ligand of both hDCIR and mDCIR1. Several different carbohydrate structures have been proposed as ligands for DCIR, suggesting that this lectin has low glycan-binding specificity. Capitalizing on the identification of LRP1 as a main ligand for hDCIR/mDCIR1, we demonstrate that these C-type lectins bind specifically to biantennary complex-type N-glycans carrying terminal galactoses and define the structural basis of such interactions for hDCIR.

LRP1 is an ubiquitous membrane receptor involved in the endocytosis, transport and lysosomal degradation of multiple ligands as well as in cell signaling.^40–43^ Over one hundred LRP1 ligands have been reported in the literature, including plasma proteins and coagulation factors, extracellular matrix proteins and proteases, cytokines, growth factors, toxins and viral proteins.^40–43^ As such, LRP1 is involved the pathogenesis of a broad range of diseases including atherosclerosis, neurodegenerative disorders, infections and cancer.^44–49^ The pathophysiological impact of LRP1 also relies on the regulatory role of this receptor in immunity.^43^ Notably, LRP1 negatively regulates inflammatory response by inhibiting TLR– mediated NF-κB activation and cytokine production in macrophages and modulates the antigen-presentation capacity of DCs to T cells.^49–60^ Given that similar anti-inflammatory functions have been attributed to human and mouse DCIR, the identification of LRP1 as a ligand for DCIR and their concomitant expression in macrophages and DCs raise the possibility that these two receptors act in concert to inhibit inflammation. However, further investigation is needed to determine whether DCIR and LRP1 interact in *cis* to regulate each other’s functions. It would also be relevant to study whether DCIR regulates apoptotic cell clearance (efferocytosis) by phagocytes, given that LRP1 is a well-known pro-efferocytosis receptor via calreticulin recognition.^58, 61–63^

Beyond LRP1, our data suggest that myeloid cells express other DCIR ligands. Indeed, although we have proven that LRP1 is a major ligand for mDCIR1, the binding of this lectin to BMDM is not completely abolished in LRP1-deficient cells. In addition, we found that mDCIR1 binds strongly to tissue neutrophils which express little to no LRP1 (data not shown).

Among other potential DCIR ligands, we recurrently identified by proteomics the cation-independent mannose-6-phosphate receptor (IGF2R), Stabilin-1 (STAB1) and macrophage mannose receptor 1 (MRC1) following affinity purification using hDCIR and/or mDCIR1 (Table S1). Similar to LRP1, these proteins are highly glycosylated endocytic cell surface receptors, indicating that DCIR binding to these proteins may require the presentation of multivalent glycans, consistent with the low affinity of CLR/carbohydrate interactions.

Among the various carbohydrate structures published as potential DCIR ligands, asialo-biantennary N-glycans have recently been reported.^9^ In agreement, we found that hDCIR and mDCIR1 interact not only with biantennary N-glycans carrying terminal galactoses but also with those bearing α-Gal epitopes. In fact, hDCIR seems to bind more to biantennary N-glycans carrying the α-Gal epitope than to those carrying single terminal β-galactoses (Figures 4C and 6A). Using X-ray crystallography and mutagenesis, we revealed the structural basis of hDCIR interaction with biantennary N-glycan carrying terminal α-Gal epitopes. We found that hDCIR interacts with the Galα1-3Galβ1-4GlcNAcβ1-2Manα via the ligation of mannose in the primary Ca^2+^ binding site of the lectin and additional interactions between amino acids and the GlcNAc and galactose residues. The requirement of galactose residues for hDCIR binding was proven to be mediated by the EPS motif, presenting weaker interaction with the mannose than a prototypical EPN motif, and the side chain of Gln226 forming hydrogen bonds with hydroxyl groups of terminal galactoses. It is notable that a similar binding mechanism was demonstrated for human BDCA-2 (CLEC4C), another CLR of the Dectin-2 cluster, that interacts with the trisaccharide Galβ1-4GlcNAcβ1-2Manα1-3 of biantennary N-glycans.^64^ In particular, the primary Ca^2+^ binding site of BDCA-2 mediates the interaction with mannoses while the side chain of BDCA-2 Gln202 (equivalent of hDCIR Gln226) forms hydrogen bonds with the hydroxyl groups of the terminal galactose residue. However, unlike DCIR, BDCA-2 has an EPN motif in its primary Ca^2+^ binding site, even though BDCA-2 interacts very weakly with non-galactosylated N-glycans. The reason for which the EPN motif of BDCA-2 is not sufficient to mediate its interaction with non-galactosylated N-glycans remains to be clarified.

The α-Gal epitope (Galα1-3Galβ1-4GlcNAc) is one of the most abundant carbohydrate antigens in all mammals except Old World monkeys, apes and humans. This epitope is also present on glycoproteins from ticks and insects. When exposed to this carbohydrate antigen, humans naturally produce anti-α-Gal antibodies which can provoke severe immune reactions including rejection of non-human organ transplants, acquired red meat allergy, a.k.a. alpha-gal syndrome, and hypersensitivity reactions to cetuximab, a chimeric mouse-human monoclonal antibody used to treat cancers of the bowel and neck.^36, 37, 65–67^ To our knowledge, hDCIR is the first human lectin described as interacting with the α-Gal epitope carried on glycoproteins, suggesting that this lectin may play an important role in α-Gal-mediated diseases. In agreement with this idea, a recent study established that DCIR on the surface of mast cells mediates the recognition and uptake of cockroach allergen, which contains the α-Gal epitope on N-glycans, and enhances cockroach/IgE-induced mast cell activation and atopic dermatitis-like inflammation.^16, 68^

Altogether, our study underscores the complexity and specificity of lectin-carbohydrate interactions within the immune system, particularly highlighting the role of DCIR in recognizing diverse glycan structures through its binding to LRP1 and potentially other glycosylated ligands. Our findings not only advance our understanding of DCIR’s functional dynamics in immune regulation but also set the stage for further investigations into its broader immunological roles and therapeutic potential.

## ACKNOWLEDGMENTS

Authors acknowledge Genotoul core facilities (ANEXPLO-IPBS and TRI-IPBS) as well as Valérie Guillet, Samuel Tranier and Lionel Mourey from the structural biophysics team of the IPBS for their help and discussions. The crystallization and macromolecular crystallography facilities used in this study are part of the Toulouse Integrated Screening Platform (PICT, IBiSA, http://www.pict.ipbs.fr). We thank the scientific staff of the European Synchrotron Radiation Facility (Grenoble, France) and ALBA (Barcelona, Spain). Authors also acknowledge Molly Migliorini for providing the GST-RAP plasmid and Audrey Hessel for further cloning and mutagenesis. Thanks to Thibaut Sanchez and Aurélien Boyancé for their help in generating the LRP1-KO cells and cell culture, respectively. Authors also acknowledge David Sancho, Anne Imberty, Etienne Meunier and Denis Hudrisier for experimental suggestions.

## AUTHOR CONTRIBUTIONS

B.B.A.R., T.S., S.R., N.G. and G.T. performed the flow cytometry experiments. S.R, A.B., M.T., F.F. and P.D. produced the recombinant lectins. B.B.A.R. and Y.R. performed the affinity purification experiments and most of biochemical analyses. T.S. and F.L. generated the LRP1-deficicent cells that were further analyzed by T.S., Y.R. and N.G. Cell culture was done by B.B.A.R., T.S., S.R., N.G, G.T. and Y.R. A.B. and Y.R. performed the Lectin ELISA. CE-ESI-MS analysis of N-glycans was done by A.L.H.E and M.W., while proteomics analyses were performed by A.S., J.M. and O.B-S. X-ray crystallography and associated analysis were performed by S.R and P.D. Y.R. wrote the manuscript with help of O.N. and B.B.A.R. O.N. and Y.R. supervised the study.

## MATERIAL AND METHODS

### Cell isolation from mouse organs and fluids

Animal experiments were approved by the regional Ethic Committee of Toulouse Biological research Federation and by the Ministry of Higher Education and Research (APAFlS#22594). Wild-type (WT) and mDCIR1-KO (*Clec4a2^−/−^*) mice were bred in an EOPS protected area (free of specific pathogens) within the animal facility of the Institute of Pharmacology and Structural Biology. Female WT and mDCIR1-KO mice aged between 8 and 12 weeks were sacrificed by cervical dislocation under anaesthesia (4% isoflurane; Vetflurane® Virbac Denmark). Blood was collected by cardiac puncture in 1.5 ml tubes containing 40 µL of 0.5 mM EDTA (Euromedex) and cells were isolated after two steps of red blood cells lysis (160 mM NH_4_Cl (Sigma); 10mM KHCO3 (Sigma) and 0.1 mM EDTA). Spleen and thymus were dissected and mechanically dissociated to obtain a single cell suspension. After lysis of red blood cells, the single cell suspension was filtered through a 70 µm cell strainer. Lungs were dissected, enzymatically digested using 100 mg/mL of collagenase D (Sigma) and 10 mg/mL of DNase I (Sigma) for 30 minutes under agitation at 37°C, and then mechanically dissociated using “lung 01.02” and “lung 02.01” programs on the gentleMACS^TM^ dissociator (Miltenyi Biotec) to obtain single cell suspension. After lysis of red blood cells, cells were filtered through a 40 µm cells strainer. Bone marrow cells were recovered from hips, femurs and tibias of hind legs by a centrifugation-based isolation method. After lysis of red blood cells, cells were filtered through a 70 µm cells strainer.

### Mouse bone marrow-derived macrophages and dendritic cells

In order to generate bone marrow-derived macrophages (BMDM), bone marrow cells (isolated as described above) were cultured in DMEM Medium 1X GlutaMax (GIBCO) with 10% of heat-inactivated fetal bovine serum (FBS; Sigma-Aldrich), 100 units/ml of penicillin and 100 µg/ml of streptomycin (Gibco), 10 mM HEPES (Gibco) and 20 ng/ml of recombinant murine M-CSF (Peprotech) in petri dishes (100 mm × 15 mm) at 37°C in a humidified atmosphere with 5% CO2 for 7-10 days. Half the medium was replaced every 2-3 days for the duration of the culture. On completion of differentiation, we removed cell supernatant and harvest BMDM (adherent) by incubating them for 10 minutes in cell detachment buffer (Dulbecco’s Phosphate Buffered Saline containing 10 mM EDTA) followed by gentle mechanical removal using cell scrapers. BMDM in PBS/EDTA were transferred in 50 ml conical tubes (Falcon) and the cell detachment buffer was neutralized by adding the same volume of DMEM with 10% of heat-inactivated FBS. A similar protocol was followed to obtain bone marrow-derived dendritic cells (BMDC), except that we used 20 ng/ml of recombinant murine GM-CSF (Peprotech or Miltenyi biotec) instead of M-CSF, that differentiation was carried out over 12 days, and that we kept the culture supernatant which contained a significant proportion of the BMDC.

### Human blood monocyte-derived macrophages and dendritic cells

Monocytes from healthy subjects were isolated from buffy coats provided by Etablissement Français du Sang, Toulouse, under contract 21/PLER/TOU/IPBS01/20130042. This contract was approved by the French Ministry of Science and Technology (agreement number AC 2009921) in compliance with articles L1243-4 and R1243-61 of the French Public Health Code. Written consent of all donors was provided before sample collection and the sex of subjects remained unknown. Monocytes from healthy subjects were isolated and differentiated into macrophages (mo-Mac) or dendritic cells (mo-DC) as previously described.^69^ Briefly, human peripheral blood mononuclear cells were recovered by gradient centrifugation on Ficoll-Paque Plus (GE Healthcare) and monocytes were further purified using CD14 microbead positive selection and MACS separation columns (Miltenyi Biotec). After purification, monocytes were allowed to adhere to glass coverslips (VWR international) in 6-well plates (Thermo Scientific) at 1.5x10^6^ cells per well for 1 h at 37°C in warm RPMI-1640 medium (GIBCO). The medium was then supplemented to a final concentration of 10% FBS (Sigma-Aldrich) and 20 ng/ml of recombinant human M-CSF (Peprotech) for differentiation in mo-Mac or 10 ng/mL recombinant human GM-CSF (Peprotech) plus 20 ng/ mL recombinant human IL-4 (Peprotech) for differentiation in mo-DC. Cells were allowed to differentiate at 37°C in a humidified atmosphere with 5% CO_2_ for at least 7 days and cell medium was renewed at day 3. At the end of the differentiation, mo-Mac (adherent) were harvested as described for BMDM, while mo-DC (non-adherent) were recovered in the cell culture supernatant.

### Generation and differentiation of HoxB8-immortalized myeloid progenitor cells

Bone marrow cells were harvested from Cas9-expressing mouse and hematopoietic progenitor cells were purified by centrifugation using Ficoll-Paque (Cytiva).^70^ Then, hematopoietic cells were pre-stimulated for 2 days with complete RPMI 1640 medium (Gibco) supplemented with 15% FCS (Pan-Biotech), 1% antibiotic-antimycotic (Gibco), 10 ng/mL recombinant mouse IL-3, 10 ng/mL recombinant mouse IL-6 and 10 ng/mL recombinant mouse SCF (Miltenyi-Biotec). Then, 2x10^5^ cells/well were seeded on a 6-well culture plate in a so-called myeloid medium containing RPMI 1640 medium, supplemented with 10% FCS, 1% antibiotic-antimycotic (Gibco), 20 ng/mL recombinant mouse GM-CSF (Miltenyi-Biotec) and 0.5 μM ß-estradiol (Sigma-Aldrich) and transduced with ER-HoxB8 retrovirus using Lentiblast Premium (OZ Biosciences).^71^ After two weeks, only immortalized ER-HoxB8 progenitors continue to proliferate. HoxB8 progenitors were then passaged every 2 days in myeloid medium. To differentiate HoxB8 progenitors into macrophages, cells were washed twice in PBS and 5x10^5^ cells were plated in one well of a 6-well plate in complete medium containing 20 ng/mL of mouse M-CSF (Miltenyi-Biotec). On day 3 and day 6, cells were fed by adding an equal amount of fresh medium. Differentiated HoxB8 macrophages were harvested by detachment for 5 minutes in PBS containing 10 mM EDTA at 37°C.

### CRISPR-Cas9-mediated gene silencing in HoxB8-Cas9 progenitor cells

For individual gene knockdown, the sgRNA corresponding to each gene was cloned into the pLenti-sgRNA vector (Addgene #71409) with BsmBI digestion (New Englands Biolabs) and sticky end ligation (New Englands Biolabs). The sgRNA sequence used to generate LRP1 knockout cells is (5’-3’): *CACC<u>G</u>ATGCCAATGAGACCGTATGC*. A luciferase (Luc) targeting sgRNA was used as a control (5’-GGCGCGGTCGGTAAAGTTGT-3’). To generate lentiviral particles, Lenti-X 293T cells (Clontech) were co-transfected with pMDL (Addgene #12251), pREV (Addgene #12253), pVSV-G (Addgene #12259), and specific sgRNA-cloned plasmids using Lipofectamine 3000 (Thermo-Fisher) and OptiMEM medium (Gibco) according to the manufacturer’s instructions for 72 hours. 2x10^5^ HoxB8 myeloid progenitor cells were transduced in 1 mL of myeloid medium containing the pLenti-sgRNA lentiviral particles using Lentiblast Premium (OZ Biosciences). After 24 hours, all cells were centrifuged for 5 minutes at 300 x g and the medium was replaced with myeloid medium containing 7 µg/mL puromycin (Invivogen) to select cells transduced for 72 hours.

### Lec2 and Lec8 cell lines

Chinese Hamster Ovary (CHO) mutant Lec2 (CRL-1736) and Lec8 (CRL-1737) cell lines were purchased from the American Type Culture Collection (ATCC) and handled upon receipt according to their instructions. Cells were cultured at 37°C in an humidified atmosphere with 5% CO2, in T75 flasks (Sarstedt # 833911002) containing MEM α with nucleosides (Gibco # 22571038). Cells were passaged every 2-3 days, with the use of trypsin-EDTA 0.05% (Gibco # 25300054) for detachment. Cells with less than 10 passages were used for the experiments.

### Silencing of LRP1 in human monocyte-derived macrophages

Targeted gene silencing in human mo-Mac was performed with a forward transfection protocol as previously described.^72^ Briefly, mo-Mac were transfected with 200 nM of ON-TARGETplus SMARTpool siRNA targeting human LRP1 (Horizon Discovery #L-004721-00-0005) or ON-TARGETplus non-targeting control pool (Horizon Discovery #D-001810-10-05) using HiPerFect (final concentration 3.0% (vol/vol); Qiagen #301707). After a six-hour transfection, RPMI 1640 medium with GlutaMAX™ and HEPES (Gibco #72400054) supplemented with FBS (final concentration 10%; Sigma-Aldrich #1943609-65-1) and recombinant human M-CSF (final concentration 20 ng/mL; Peprotech #300-25) was added to each well, and the cells were allowed to recuperate. Mo-Mac were used for lectin binding assays at day 3 post-transfection.

Assessment of the efficiency of gene silencing was done after three days post-transfection by real-time quantitative PCR (RT-qPCR). Mo-Mac were lysed using RLT buffer (Qiagen #79216) supplemented with 1% β-mercaptoethanol and stored at -70°C. Total RNA was extracted from the RLT samples using the RNEasy mini kit (Qiagen #74104). RNA amount and purity (absorbances at 260/230 nm and 260/280 nm) were measured using a NanoDrop spectrophotometer. RNA was then reverse transcribed from 200 ng total RNA with Moloney murine leukemia virus reverse transcriptase (Invitrogen #28025013) using oligo (dT)_18_ (Thermo Scientific #SO131) for priming according to the manufacturer’s protocol. Specific qPCR primers for human *LRP1* were obtained from OriGene (#HP206040). Specific qPCR primers for human *GAPDH* were designed using Primer-BLAST (https://www.ncbi.nlm.nih.gov/tools/primer-blast/) (forward primer (5’-3’): TCGGAGTCAACGGATTTGGTC ; reverse primer (5’-3’): GACGGTGCCATGGAATTTGC) and purchased from Sigma-Aldrich. RT-qPCR was performed with gene-targeted primers using TB Green Premix Ex Taq II (Tli RNase H Plus) (Takara #RR82WR) according to the manufacturer’s protocol. All RT-qPCR reactions were carried out using a 7500 Real-Time PCR System and data were analyzed using the 7500 Software version 2.3 (Applied Biosystems). RT-qPCR data are reported as the *LRP1* transcript abundance in each sample compared to the *GAPDH* housekeeping gene as calculated by the comparative cycle threshold method (2^(-ΔCt^)), and normalized to the non-transfected mo-Mac samples (2^(-ΔΔCt^)).

### Production and biotinylation of recombinant lectins

Across the study, we used lectins that have been biotinylated chemically (Figures 1, 2B-D and 3A-C) or enzymatically (Figures 2A, 2E, 3D and 4C-D and 6). For chemically labeled biotinylated proteins, hDCIR-ECD (Uniprot Q9UMR7-1, amino acids 70-237, CCDS8590.1), mDCIR1-ECD (Uniprot Q923C7, amino acids 70 to 262) and mDCIR1-τιECD (E221A) with N-terminal His-Tag and TEV proteolytic site (HHHHHHIEGRGGGGG) were produced and purified as described.^73^ Chemical biotinylation of lectin ECD was performed using EZ-Link™ Sulfo-NHS-LC-Biotin (Thermo Scientific #21335) according to manufacturer instructions. Follwing biotinylation, excess of biotin was removed using Zeba Spin Desalting Columns (Thermo Scientific). For Enzymatically labeled biotinylated proteins, the coding sequences of mDCIR1-ECD (Uniprot Q9QZ15, amino acids 70-238, CCDS20510.1) and hDCIR-ECD (Uniprot Q9UMR7-1, amino acids 70-237, CCDS8590.1) with a N-terminal Avi-tag (GLNDIFEAQKIEWHE) were obtained as synthetic genes (Eurofins Genomics) in the pJ411 expression plasmid (plasmid with high copy number, kanamycine resistance; kindly gifted by Dr. Guillaume Ferré). For the lectin mutants, site-directed mutagenesis was performed by PCR, and the sequences were verified by Sanger sequencing (Eurofins Genomics). hDCIR-ECD and mDCIR1-ECD were expressed in *E. coli* BL21 Star™ (DE3) cells (Invitrogen) in 500 mL of 2YT medium supplemented with 50 μg/mL kanamycin at 37°C and 160 rpm. When the culture had reached an OD_600_ of 0.8-1.0, protein expression was induced for 4 hours by addition of 1 mM isopropyl-β-D-thiogalctopyranoside (IPTG). In the transformed bacteria, the production of recombinant lectins is done in insoluble form as inclusion bodies. After a centrifugation at 5,000 x g for 10 minutes, the cell pellets were resuspended in 50 mL of ice-cold lysis buffer (50 mM Tris-HCl [pH 8.0], 100 mM NaCl, 1.0% Triton X-100, 10 mM ethylenediaminetetraacetic acid (EDTA), supplemented with Benzonase Safety Plus Emprove Expert (0.5 µL/g of wet cell pellet; Merck, #1037731010), and mechanically disrupted using an Avestin Emulsiflex-C5 (3 passes at 500-1000 bar). The insoluble fraction of disrupted cells was then washed three times, successively with 50 mL of wash buffer 1 (50 mM Tris-HCl pH 8.0, 500 mM NaCl, 1 % Triton X-100, 10 mM EDTA, 5 mM dithiothreitol (DTT), 2 M urea), 50 mL of wash buffer 2 (50 mM Tris-HCl pH 8.0, 100 mM NaCl, 10 mM DTT), and 50 mL of wash buffer 3 (50 mM Tris-HCl pH 8.0, 100 mM NaCl), with in-between centrifugations of 30 minutes at 15,000 x g and 4°C. The inclusion bodies were solubilized in 10 mL of denaturing buffer (50 mM 2-(*N*-morpholino)ethanesulfonic acid pH 6.5, 100 mM NaCl, 10 mM EDTA, 10 mM β-mercaptoethanol, 6 M guanidine-HCl) overnight at 4°C on a rotating wheel. The solubilized protein (whose amount is usually around 50 mg at this step, as measured at 280 nm using a NanoDrop spectrophotometer) was slowly diluted drop by drop using a peristaltic pump into 500 mL of refolding buffer (50 mM Tris-HCl pH 8.5, 100 mM NaCl, 0.4 M L-arginine, 5 mM CaCl_2_, 10 mM reduced glutathione, 1 mM oxidized glutathione, 0.1 mM phenylmethylsulfonyl fluoride) at 4°C, and left to refold overnight at 4°C under gentle stirring. The diluted solution was dialyzed in Spectra/Por™ 3 RC Dialysis Membrane Tubing 3.5 kDa MWCO (Spectrum Laboratories #132724) against 5L of dialysis buffer (20 mM 4-(2-hydroxyethyl)-1-piperazineethanesulfonic acid (HEPES) pH 7.5, 100 mM NaCl, 5 mM CaCl_2_) 5 times over two days at 4°C. Aggregates were removed by centrifugation at 15,000 x g for 30 minutes at 4°C, and filtration on a 0.2 µm polyethersulfone membrane (VWR, #514-0340). The solution was then concentrated by ultrafiltration, using successively an Ultracel regenerated cellulose disk (molecular weight cutoff, 10 kDa ; Millipore, #PLGC06210) mounted on a stirred cell (Amicon, #UFSC20001), and a Vivaspin 20 concentrator (molecular weight cutoff (MWCO) 10 kDa ; Sartorius, #VS2002). The proteins were then purified by size-exclusion chromatography using a HiLoad 16/600 Superdex 75 pg Prep grade column (Cytiva #28989333) equilibrated with gel filtration buffer (20 mM HEPES pH 7.5, 100 mM NaCl, 5 mM CaCl_2_). The purity of the fractions of interest was verified by SDS-PAGE and Quick Blue Protein Stain (LubioScience #LU001000). After the size-exclusion chromatography, the lectins were concentrated up to 1-5 mg/mL using a Vivaspin sample concentrators (MWCO 10 kDa), and enzymatically biotinylated using BirA ligase (Sigma-Aldrich #CS0008) overnight at 4°C according to the manufacturer’s protocol (BirA : protein ratio (w/w) of 1 : 200, molar ratio of protein : biotin of 1 : 1.5, 5 mM ATP, 5 mM MgSO_4_). The final protein concentration was determined using DC Protein Assay (Bio-Rad #5000112). Usually, starting from 500 mL bacterial culture, the final yield is 15-25 mg of recombinant lectin. The efficiency of biotinylation was verified using a streptavidin-shift assay and intact protein liquid-chromatography electrospray mass spectrometry (LC-ESI-MS).^74^

### Flow Cytometry

In order to characterize cells expressing mDCIR1 ligand(s), immune cells extracted from mouse blood and tissues as described above were first stained with 40 µg/mL of biotinylated mDCIR1-ECD or mDCIR1-ΛιECD pre-conjugated with with ¼ molar equivalents of Streptavidin-Phycoerythrin (streptavidin-PE; Biolegend) in DPBS containing Ca^2+^ and Mg^2+^ (Gibco). Cells were then blocked with mouse TruStain FcX^TM^ (Biolegend) and further stained with the LIVE/DEAD^TM^ Fixable Aqua Dead Cell Stain Kit (ThermoFisher Scientific) and the antibodies listed in Table S4. To confirm the expression pattern of mDCIR1, we used anti-mDCIR1 (clone TKKT-1; gift from Naoki Matsumoto, Japan) coupled in-house to Alexa Fluor 647 using Alexa Fluor® 647 Protein Labeling Kit (Thermo Fisher Scientific #A20173). Multicolor analysis was performed on a BD LSRFortessa^TM^ cell analyzer, and data were analyzed with the Flowjo software (FlowJo LLC, version 10.2.6). Gating strategies for analysis of immune cell populations are shown in Figure S1. Regarding lectin binding experiments with macrophages, dendritic cells and CHO cell lines, 20 μg/mL of biotinylated Lectin-ECD in DPBS containing Ca^2+^ and Mg^2+^ (Gibco) were pre-conjugates with ¼ molar equivalents of streptavidin-PE (Biolegend) in the presence or absence of 10 mM chelating agents (*i.e*., EGTA or EDTA).

### mDCIR1-ECD and hDCIR-ECD pulldown experiments and proteomic quantification

Human and murine cells were incubated in lysis buffer (20 mM Tris-HCl, 150 mM NaCl, 1 % NP-40, 5 % glycerol) containing 1X protease inhibitor cocktail (Roche) on ice, with frequent vortexing for 30 min. The cells were then lysed by 15 back and forth passes through a 23G needle followed by centrifugation at 20,000 x g at 4°C for 10 minutes to remove cellular debris. Protein concentration was measured using the DC protein assay (Bio-Rad). Ten µg of biotinylated mDCIR1-ECD, mDCIR1-ΔECD or hDCIR-ECD was incubated with Streptavidin Dynabeads (50 ul solution pre-washed 3 times with PBS pH 7.4; Thermo Fisher Scientific #65601) for 2 hours at 4°C on a rotating wheel. The unbound fraction was collected using a magnetic rack and the conjugated beads were then incubated with 1 mg of protein lysate (in 300 μl of lysis buffer) overnight at 4°C on a rotating wheel. As negative controls, 100 mM EGTA was added during the overnight incubation of DCIR-bound beads with protein lysates. The flow through (unbound proteins) was collected and the beads were washed 5 times with lysis buffer using a magnetic rack. Bound proteins were eluted by the addition of 100 mM EGTA in PBS and incubation with shaking (600 rpm) for 5 min at room temperature. The eluted proteins were cleaned up by a short SDS-PAGE run on 12% gel (i.e., until proteins migrated into the resolving gel). Proteins were then stained using quick Coomassie stain (Cliniscience #NB-45-00078-1L) and protein bands were excised using a scalpel in order to perform an in-gel trypsin digestion as described.^75^

### Proteomic quantification

Regarding proteomics of mo-DC-derived samples, tryptic peptides were resuspended in 22µl of 2% acetonitrile and 0.05% trifluoroacetic acid and analyzed by nano-liquid chromatography (LC) coupled to tandem MS, using an UltiMate 3000 system (NCS-3500RS Nano/Cap System; Thermo Fisher Scientific) coupled to an Orbitrap Q-HFX mass spectrometer (Thermo Fisher Scientific). 5µl of each sample was loaded on a C18 precolumn (300 µm inner diameter × 5 mm, Thermo Fisher Scientific) in a solvent made of 2% acetonitrile and 0.05% trifluoroacetic acid, at a flow rate of 20 µl/min. After 3 min of desalting, the precolumn was switched online with the analytical C18 column (75 µm inner diameter × 50 cm, Acclaim PepMap 2µm C18 Thermo Fisher Scientific) equilibrated in 95% solvent A (5% acetonitrile, 0.2% formic acid) and 5% solvent B (80% acetonitrile, 0.2% formic acid). Peptides were eluted using a 10%-45% gradient of solvent B over 120 min at a flow rate of 350 nl/min. The Orbitrap Q-HFX was operated in data-dependent acquisition mode with the Xcalibur software. MS survey scans were acquired with a resolution of 60,000 and an AGC target of 3e6. The 12 most intense ions were selected for fragmentation by high-energy collision induced dissociation, and the resulting fragments were analyzed at a resolution of 15000, using an AGC target of 1e5 and a maximum fill time of 54ms. Dynamic exclusion was used within 30 s to prevent repetitive selection of the same peptide.

For proteomics of mo-Mac-and BMDM-derived samples, Tryptic peptides were resuspended in 22µl of 2% acetonitrile and 0.05% trifluoroacetic acid and analyzed by nano-liquid chromatography (LC) coupled to tandem MS, using an UltiMate 3000 system (NCS-3500RS Nano/Cap System; Thermo Fisher Scientific) coupled to an Orbitrap Fusion mass spectrometer (Thermo Fisher Scientific). 5µl of each sample was loaded on a C18 precolumn (300 µm inner diameter × 5 mm, Thermo Fisher Scientific) in a solvent made of 2% acetonitrile and 0.05% trifluoroacetic acid, at a flow rate of 20 µl/min. After 5 min of desalting, the precolumn was switched online with the analytical C18 column (75 μm inner diameter × 50 cm, in-house packed with Reprosil C18) equilibrated in 95% solvent A (5% acetonitrile, 0.2% formic acid) and 5% solvent B (80% acetonitrile, 0.2% formic acid). Peptides were eluted using a 5%-50% gradient of solvent B over 110 min at a flow rate of 300 nl/min. The OrbiTrap Fusion was operated in Data Dependent Acquisition mode to automatically switch between full scan MS and MS/MS acquisition using Xcalibur software. Survey scan MS was acquired in the Orbitrap over the m/z 300–2000 range, with the resolution set to a value of 120 000 (m/z 400). The most intense ions per survey scan were selected for higher-energy collisional dissociation fragmentation (time between Master scans: 3 s), and the resulting fragments were analysed in the linear ion trap. Dynamic exclusion was employed within 60 s to prevent repetitive selection of the same peptide.

Raw MS files were processed with the Mascot software (version 2.7.0) for database search and Proline for label-free quantitative analysis (version 2.1.2).^76^ Data were searched against Mus musculus protein database (SwissProt 20180213, 16,965 sequences) or Human protein database (SwissProt 20180213, 20,259 sequences). Carbamidomethylation of cysteines was set as a fixed modification, whereas Acetyl (Protein N-term); Oxidation (M); HexNAc (N); HexNAc (S); HexNAc (T) were set as variable modifications. Specificity of trypsin/P digestion was set for cleavage after K or R, and two missed trypsin cleavage sites were allowed. The mass tolerance was set to 5 ppm for the precursor and to 20 mmu in tandem MS mode. Minimum peptide length was set to 7 amino acids, and identification results were further validated in Proline by the target decoy approach using a reverse database at both a PSM and protein false-discovery rate of 1%. For label-free relative quantification of the proteins across biological replicates and conditions, cross-assignment of peptide ions peaks was enabled inside group with a match time window of 1 min, after alignment of the runs with a tolerance of +/- 600s.

Median Ratio Fitting computes a matrix of abundance ratios calculated between any two runs from ion abundances for each protein. For each pair-wise ratio, the median of the ion ratios is then calculated and used to represent the protein ratio between these two runs. A least-squares regression is then performed to approximate the relative abundance of the protein in each run in the dataset. This abundance is finally rescaled to the sum of the ion abundances across runs. A paired student T-test was then performed on log10 transformed values to analyze differences in protein abundance in all biologic group comparisons. Significance level was set at p-value =0.05, and ratios were considered relevant if higher than +/- 5.

### Immunoprecipitation of mouse LRP1 and human LRP1

Human and murine cells were lysed as described previously in the previous section. After protein quantification (DC protein assay; Bio-Rad), 1 mg of protein lysate (in 300 μl of lysis buffer) from BMDM, BMDC, mo-DC or mo-Mac were pre-cleared by incubating with Protein G Dynabeads (50 μl of Protein G Dynabeads slurry pre-washed 3 times with PBS containing 0.02 % Tween-20) for 1 h at room temperature on a rotating wheel to remove any non-specific interacting proteins. The unbound fraction (cleared lysates) was collected using a magnetic rack. Antibodies were coupled to Dynabeads G proteins after 3 washes of 50 μl Dynabeads G protein slurry with PBS containing 0.02 % Tween-20 and incubation for 2 hours at 4°C on a rotating wheel with 100 µl PBS containing 0.02 % Tween-20 and 10 µg anti-LRP1 (Abcam, #ab92544) or isotype control (Abcam, #ab172730). After supernatant removal using a magnetic rack, cleared lysates were incubated with antibody/protein G Dynabeads overnight at 4 °C on a rotating wheel. The unbound fraction was collected and beads were washed 4 times with 500 µl of lysis buffer using a magnetic rack. Bound proteins were eluted in 50 µl of 1x Laemmli Sample Buffer (Bio-Rad) at 95°C for 5 minutes, collected using a magnetic rack and finally frozen. For PNGase treatment of LRP1, bound proteins were eluted with 40 µl of 2x SDS at 95°C for 5 minutes and collected using a magnetic rack and finally dried using a vacuum concentrator. Dried proteins were resolubilized in 10 µL of Tris-HCl buffer at pH 7 and 20 µl of MQ water. Subsequently, 20 µL of 1U PNGaseF (Roche) in 2.5X PBS containing 2% NP-40 was added and the mixture prior to overnight incubation at 37°C.

### SDS-PAGE, Western blot and Lectin blot

Protein lysates were separated using 4-12% Bis-Tris SDS-polyacrylamide gel electrophoresis (SDS-PAGE) (Thermo Fisher Scientific, #NP0321) for 1-2 h at 150V, followed by electroblotting onto 0.2 μm nitrocellulose or pvdf membranes (Bio-Rad, #1704270 or 1704272) using the Trans-Blot Turbo Transfer System. Precision Plus Protein™ Kaleidoscope™ Prestained Protein was used as protein ladder standards (Bio-Rad #1610375). For high molecular weight proteins including LRP1 α-chain, cell lysates were separated using non-reducing 6% SDS-PAGE (home-made) at 4°C starting at 50V for 1 h, 70V for 1 h and then 80V for 3 h or until the 75 k Da marker migrated completely off the gel. Following electrophoresis, gels were either fixed in quick Coomassie stain (Cliniscience #NB-45-00078-1L), or proteins were electroblotted onto 0.45 μM PVDF membranes (Thermo Scientific #88585) at 30V overnight at 4 °C. Membranes were blocked in 5% BSA (Serva, #11932 or Roche #03117057001) in PBS containing 0.05 % Tween-20 (Sigma-Aldrich #P-1379) for 1 h at room temperature, followed by incubation with one of the following primary antibodies overnight at 4°C: anti-LRP1 (Abcam, #ab92544), anti-LRP1 (Sigma-Aldrich, #L2295), anti-LRPAP1 (Abcam, #ab76500), anti-actin (Cell Signaling Technology, #4967), and anti-alpha-gal epitope (Absolute antibody # Ab00532-10.0 HRP) . Following extensive washing in TBS containing 0.05% Tween-20, membranes were incubated with one of the following HRP-conjugated secondary antibodies for 1 h at room temperature: anti-rabbit HRP (Advansta, #R-05072-500), anti-mouse HRP (Advansta, #R-05071-500), anti-rat HRP (Abcam, #ab97057). Membranes were developed using WesternBright Quantum HRP substrate (Advansta, #K-12042). For Lectin blots, PVDF membranes were blocked in 5% BSA in TSM buffer (20 mM Tris, pH 7.4, 150 mM NaCl, 1 mM CaCl2 and 2 mM MgCl2) containing 0.05% Tween-20 for 1 h at room temperature. Following this, 3-5 µg/mL of mDCIR1-ECD or mDCIR1-ΔECD (pre-conjugated with ¼ molar equivalent of streptavidin-HRP; Thermo Fisher Scientific #21124) diluted in TSM was incubated for 1 h at room temperature. After extensive washing in TSM containing 0.05% Tween-20, membranes were developed using WesternBright Quantum HRP substrate (Advansta).

### Isolation of LRP1 and glycomic analysis

Purification of mouse macrophage LRP1 α-chain was performed by immunoprecipitation, as described in the previous section, using 3 mg of BMDM protein lysate. Affinity purification of LRP α-chain from human serum was performed as previously described.^77^ Briefly, 800 ml of human Serum (Sigma Aldrich # H4522) were thawed at 37°C after addition of cOmplete™, EDTA-free Protease Inhibitor Cocktail (2 tablets/100 ml of serum; Roche #11873580001) and 8 ml of 500 mM CaCl2. Serum was dialyzed in Spectra/Por™ 2 RC Dialysis Membrane Tubing 12-14 kDa MWCO (Spectrum Laboratories #132680) against 4 L of 50 mM Tris-HCl, 150 mM NaCl, pH 7.5, with 1 mM CaCl2 and 1 mM MgCl2 for 12 h at 4°C. The dialyzed serum was subjected to 2 x centrifugation at 10,000 g for 10 min at 4°C and then filtered on a 0.45 μm Sterile Disposable Bottle Top Filters (Thermo Scientific Nalgene #295-4545). An affinity matrix was prepared by coupling 50 mg of His-tagged GST-RAP to 20 ml of NHS-activated Sepharose 4 Fast Flow (GE Healthcare), according to the manufacturer’s instructions. Serum was then incubated with the affinity matrix overnight at 4°C on a rotating wheel. The mixture (serum and GST-RAP coupled NHS-activated sepharose) was loaded to an empty glass chromatography column (Bio-Rad #7372512) to remove the unbound proteins (flow through) and the Sepharose medium was washed 6 times with 50 mL of 0.1 M Tris-HCl, pH 8.5. The bound proteins were eluted from the sepharose medium with 7 x 15 mL of elution buffer (0.1 M sodium acetate, 0.5 M NaCl pH 4.5) and the elution fractions were rapidly neutralized by the addition of 2M Tris-base (Sigma Aldrich). Elution fractions were loaded on a SDS-PAGE and presence of LRP1 α-chain revealed by far western blot using His-tagged GST-LRPAP and a secondary anti-His antibody coupled to HRP. Elution fractions containing LRP1 α-chain were pooled and concentrated using Amicon Ultra-15 30 kDa (Merck #UCF903024). Aliquots of purified LRP1 α-chains (from mouse macrophage and human serum) were separated by SDS-PAGE, gel bands corresponding to the LRP1 protein were excised and the *N*-glycans were attained from an overnight in-gel *N*-glycan release with 2U PNGaseF (Roche) at 37°C. Capillary electrophoresis-electrospray ionization-mass spectrometry (CE-ESI-MS) analysis of LRP1 total released *N*-glycans with sialic acid linkage-specific derivatization and reducing end labeling was performed as described previously.^35^ Briefly, sialic acids were derivatized by ethyl esterification (α2,6-linked sialic acids) and subsequent ammonia amidation (α2,3-linked sialic acids) by incubation in 20 µL ethyl esterification reagent (250 mM EDC and 250 mM HOBt in ethanol) for 30 min at 37°C, followed by amidation with 4 µL 28% NH4OH for 30 min at 37°C.

Derivatized *N*-glycans were purified by cotton-hydrophilic interaction liquid chromatography (HILIC) solid phase extraction (SPE) and dried by vacuum centrifugation. Next, 2 μL Girard’s reagent P (GirP) (50 mM GirP in 90% ethanol and 10% glacial acetic acid (HAc) was added and incubated for 1h at 60°C. The glycans were dried by vacuum concentration and resuspended in 6 µL leading electrolyte (400 mM ammonium acetate pH = 3.2) before analysis. CE-ESI-MS analyses were performed on a CESI 8000 system (Sciex, Farmingham, MA, USA) using a neutrally coated capillary (91 cm long, 30 μm i.d., 150 μm o.d.; SCIEX) coupled to an impact HD quadrupole–time-of-flight mass spectrometer (Bruker Daltonics) utilizing dopant enriched nitrogen (DEN)-gas supply (nanoBooster technology from Bruker Daltonics). Separation was conducted at 20 °C using a background electrolyte (BGE) consisting of 20% HAc. The samples were loaded onto the capillary by hydrodynamic injection of 1 psi for 60 s (8.7 nL/1.35% capillary volume). The MS was operated positive-ionization mode using the following parameters: capillary voltage -1000 V, drying gas temperature 150 °C, drying gas flow rate 1.2 L/min. MS as well as tandem MS (MS/MS) spectra were acquired with mass range *m/z* = 200 to *m/z* = 2000 using 1 Hz as the spectral acquisition rate. MS/MS was performed on the three most abundant precursor ions. The collision energies were set as a linear curve in a *m/z* dependent manner, ranging from 55 eV at *m/z* 700 to 124 eV at *m/z* 1800 for all charge states (1–5), applying a basic stepping mode. Glycans compositions were identified based on their exact mass, migration order, fragmentation spectra and previous described structures in literature using DataAnalysis software version 5.0 (Bruker Daltonics). Extracted ion electropherograms (EIEs) were generated for the first three isotopes of the singly, doubly, and triply charged analytes with a width of ±0.015 *m/z* units and the area of the EIEs were integrated. The obtained areas were normalized to 100%, whereafter the glycovariants were relatively.

### Glycan coupling to BSA

For coupling glycan to BSA, the latter was dissolved at 2 mg/ml in PBS pH 7.2 containing 5 mM EDTA and reduced by adding a 20 molar equivalents of tris(2-carboxyéthyl)phosphine (TCEP) followed by incubation for 30 minutes at room temperature. Excess of TCEP was then removed by size-exclusion chromatography using a Zeba Spin Desalting Column (Thermo Scientific) pre-washed 4 times with PBS pH 7.2 containg 5 mM EDTA and the concentration of reduced BSA was measured using a NanoDrop spectrophotometer. A 25-molar equivalent of glycan-maleimide was added to the reduced BSA and the mixture was incubated under agitation for 2 hours at room temperature. The excess of glycan-maleimide was removed using a Zeba Spin Desalting Column pre-washed 4 times with PBS pH 7.2 and the concentration of BSA-glycan was assessed using a NanoDrop spectrophotometer. The relative proportion of glycosylation was assessed by intact protein LC-MS. On average, we found more than 3 carbohydrate units per BSA for all N-glycans, with the exception of sialylated N-glycans, for which we obtained an average of 1.31 carbohydrate units per BSA.

### Lectin ELISA

Nunc MaxiSorp 96-well plates (Thermo Scientific # 439454) were coated with 100 μl/well of BSA-glycans (concentration ranging from 0.1 to 10 μg/ml) diluted in TBS (20 mM Tris-HCl, 150 mM NaCl pH 7.4) overnight at 4°C. Wells were then blocked with 250 μl/well of 1% BSA in TSM buffer (20 mM Tris-HCl, pH 7.4, 150 mM NaCl, 1 mM CaCl_2_ and 2 mM MgCl_2_) for 1 hour at room temperature. After 3 washings with 250 μl/well of TSM containing 0.05% Tween, 100 μl/well, 10 μg/ml of biotinylated DCIR-ECD dissolved in TSM (and pre-incubated for 15 minutes at room temperature with ¼ molar equivalent of streptavidin-HRP; Thermo Scientific #21124) were added followed by incubation for 3 hours at room temperature. Wells were then washed 5 times with 250 μl/well of TSM containing 0.05% Tween before the addition of 100 μl/well of TMB solution (3,3’,5,5’-Tetramethylbenzidine; Invitrogen). After few minutes, the reaction was stop by adding 50 μl/well of 2N H_2_SO_4_. The absorbance was immediately measured at 450 nm and 570 nm (for wavelength correction) using a CLARIOstar Plus microplate reader (BMG Labtech).

### Intact protein LC-ESI-MS of conjugated DCIR and BSA

NanoLC-MS analyses were performed with 5 μL of purified protein diluted in loading buffer (2% acetonitrile (ACN), 0.1% trifluoro acetic acid (TFA)) to a final concentration of 1 μM, by employing a nanoRS UHPLC system (Dionex), coupled to a Fusion Tribrid Orbitrap mass spectrometer (Thermo Fisher Scientific). Samples were loaded onto a reverse-phase C4 pre-column (300 μm i.d. x 5mm; Thermo Fisher Scientific) at 20 μL/min in 2% ACN and 0.05% TFA. After 5 min of desalting, the precolumn was switched online to a home-made C4 analytical nanocolumn (75 μm i.d. x 15 cm) packed with C4 Reprosil (Cluzeau CIL), equilibrated in 95% solvent A (0.2% formic acid (FA)) and 5% solvent B (0.2% FA in ACN). Proteins were eluted using a linear gradient from 5% to 30% B in 38 min for DCIR and 15% to 50% B in 21 min for BSA, at 300 nL/min. MS scans were acquired in positive mode in the 700-2,000 m/z range for DCIR and 500-2,000 m/z range for BSA with a resolution set at 7,500. The DCIR spectra were deconvoluted and semi-quantified with Unidec 4.4.0 using the following parameters: m/z range: 700–2,000 Th; no background subtraction, no smoothing; charge range: 10–30; mass range: 10,000–30,000 Da; sample mass: every 1 Da; peak detection range: 50 Da, and peak detection threshold: 0.05.^78^ The BSA spectra were deconvoluted and semi-quantified with Unidec 4.4.0 using the following parameters: m/z range: 700–2,000 Th; subtract Curved: 10, Gaussian smoothing: 5; charge range: 30–70; mass range: 50,000–90,000 Da; sample mass: every 1 Da; peak detection range: 200 Da, and peak detection threshold: 0.05.

### **X-** ray crystallography and analysis

For the purpose of X-ray crystallography experiments, recombinant hDCIR-ECD devoid of tag was used ; the sequence encoding the N-terminal Avi-tag was removed from the expression plasmid using In-Fusion® cloning (Takara #638948) prior to protein expression and purification. All crystallization experiments were carried out using the sitting drop vapor diffusion method. The initial conditions were screened using the JCSG Plus screen (Molecular Dimensions) on a 96-well 2-drop crystallization plate (Molecular Dimensions) at 12°C. The protein concentration for crystallization assays was 5 to 10 mg/ml in 20 mM HEPES pH 7.5, 100mM NaCl, 5mM CaCl_2_. Co-crystallization of the hDCIR/carbohydrate ligand complex was carried out under the conditions of 0.1 M Tris pH 8.5, 20% w/v PEG 8000, 0.2 M MgCl_2_ (227.39 μM of hDCIR ECD and 20 mM of ligand). The crystals were cryo-chilled directly in liquid nitrogen. X-ray diffraction data sets were collected at ESRF or Alba using beamline ID30b or XALOC respectively. The data were processed using XDS and Phenix.^79, 80^ The structures were solved by molecular replacement using the X-ray structure of the CRD domain of hDCIR (PDB code 5b1w or 5b1x) for the hDCIR/glycan complex. The CHARMM-GUI website (http://www.charmm-gui.org) was used to generate a model of the biantennary complex type N-glycan containing terminal α-Gal epitopes. Several iterative cycles of model building in Coot and refinement in Phenix were performed until convergence was reached.^81, 82^ Crystallography figures were generated using PyMol Version 2.3.0 (Schrödinger, LLC). The 2D interaction maps were designed based on Ligplot+ v.2.2 (https://www.ebi.ac.uk/thornton-srv/sofware/LigPlus/).

## QUANTIFICATION AND STATISTICAL ANALYSIS

Statistical details of experiments are indicated in the figure legends or method details section. All data analyses were performed with Prism software (GraphPad). All data are presented as mean ± standard deviation (SD), and error bars represent the SD of at least three biological replicates, unless otherwise indicated in the figure legends. No statistical analysis was used to predetermine sample size.

## SUPPLEMENTAL INFORMATION

Supplemental figures and tables can be found in separate files.

## REFERENCES

1. Sousa, C.R. e, Yamasaki, S., and Brown, G.D. (2024). Myeloid C-type lectin receptors in innate immune recognition. Immunity 57, 700–717. 10.1016/j.immuni.2024.03.005.

2. Brown, G.D., Willment, J.A., and Whitehead, L. (2018). C-type lectins in immunity and homeostasis. Nat Rev Immunol 18, 374–389. 10.1038/s41577-018-0004-8.

3. Kaifu, T., and Iwakura, Y. (2016). Dendritic Cell Immunoreceptor (DCIR): An ITIM-Harboring C-Type Lectin Receptor. In C-Type Lectin Receptors in Immunity., pp. 101–113. 10.1007/978-4-431-56015-9_7.

4. Dudziak, D., Kamphorst, A.O., Heidkamp, G.F., Buchholz, V.R., Trumpfheller, C., Yamazaki, S., Cheong, C., Liu, K., Lee, H.-W., Park, C.G., et al. (2007). Differential Antigen Processing by Dendritic Cell Subsets in Vivo. Science 315, 107–111. 10.1126/science.1136080.

5. Ishiguro, T., Fukawa, T., Akaki, K., Nagaoka, K., Takeda, T., Iwakura, Y., Inaba, K., and Takahara, K. (2017). Absence of DCIR1 reduces the mortality rate of endotoxemic hepatitis in mice. European journal of immunology 47, 704–712. 10.1002/eji.201646814.

6. Sun, H., Tang, C., Chung, S.-H., Ye, X.-Q., Makusheva, Y., Han, W., Kubo, M., Shichino, S., Ueha, S., Matsushima, K., et al. (2022). Blocking DCIR mitigates colitis and prevents colorectal tumors by enhancing the GM-CSF-STAT5 pathway. Cell Reports 40, 111158. 10.1016/j.celrep.2022.111158.

7. Stoff, M., Ebbecke, T., Ciurkiewicz, M., Pavasutthipaisit, S., Mayer-Lambertz, S., Störk, T., Pavelko, K.D., Baumgärtner, W., Jung, K., Lepenies, B., et al. (2021). C-type lectin receptor DCIR contributes to hippocampal injury in acute neurotropic virus infection. Sci Rep-uk 11, 23819. 10.1038/s41598-021-03201-2.

8. Troegeler, A., Mercier, I., Cougoule, C., Pietretti, D., Colom, A., Duval, C., Manh, T.-P.V., Capilla, F., Poincloux, R., Pingris, K., et al. (2017). C-type lectin receptor DCIR modulates immunity to tuberculosis by sustaining type I interferon signaling in dendritic cells. Proc National Acad Sci 114, E540–E549. 10.1073/pnas.1613254114.

9. Kaifu, T., Yabe, R., Maruhashi, T., Chung, S.-H., Tateno, H., Fujikado, N., Hirabayashi, J., and Iwakura, Y. (2021). DCIR and its ligand asialo-biantennary N-glycan regulate DC function and osteoclastogenesis. J Exp Medicine 218, e20210435. 10.1084/jem.20210435.

10. Fujikado, N., Saijo, S., Yonezawa, T., Shimamori, K., Ishii, A., Sugai, S., Kotaki, H., Sudo, K., Nose, M., and Iwakura, Y. (2008). Dcir deficiency causes development of autoimmune diseases in mice due to excess expansion of dendritic cells. Nat Med 14, 176–180. 10.1038/nm1697.

11. Maruhashi, T., Kaifu, T., Yabe, R., Seno, A., Chung, S.-H., Fujikado, N., and Iwakura, Y. (2015). DCIR maintains bone homeostasis by regulating IFN-γ production in T cells. J Immunol 194, 5681–5691. 10.4049/jimmunol.1500273.

12. Lambert, A.A., Barabé, F., Gilbert, C., and Tremblay, M.J. (2011). DCIR-mediated enhancement of HIV-1 infection requires the ITIM-associated signal transduction pathway. Blood 117, 6589–6599. 10.1182/blood-2011-01-331363.

13. Maglinao, M., Klopfleisch, R., Seeberger, P.H., and Lepenies, B. (2013). The C-type lectin receptor DCIR is crucial for the development of experimental cerebral malaria. Journal of immunology (Baltimore, Md. : 1950) *191*, 2551–2559. 10.4049/jimmunol.1203451.

14. Hütter, J., Eriksson, M., Johannssen, T., Klopfleisch, R., Smolinski, D. von, Gruber, A.D., Seeberger, P.H., and Lepenies, B. (2014). Role of the C-type lectin receptors MCL and DCIR in experimental colitis. PloS one 9, e103281. 10.1371/journal.pone.0103281.

15. Seno, A., Maruhashi, T., Kaifu, T., Yabe, R., Fujikado, N., Ma, G., Ikarashi, T., Kakuta, S., and Iwakura, Y. (2015). Exacerbation of experimental autoimmune encephalomyelitis in mice deficient for DCIR, an inhibitory C-type lectin receptor. Experimental animals / Japanese Association for Laboratory Animal Science 64, 109–119. 10.1538/expanim.14-0079.

16. Luo, X., Chen, J., Yang, H., Hu, X., Alphonse, M.P., Shen, Y., Kawakami, Y., Zhou, X., Tu, W., Kawakami, T., et al. (2022). Dendritic cell immunoreceptor drives atopic dermatitis by modulating oxidized CaMKII-involved mast cell activation. Jci Insight 7, e152559. 10.1172/jci.insight.152559.

17. Lambert, A.A., Azzi, A., Lin, S.-X., Allaire, G., St-Gelais, K.P., Tremblay, M.J., and Gilbert, C. (2013). Dendritic cell immunoreceptor is a new target for anti-AIDS drug development: identification of DCIR/HIV-1 inhibitors. PloS one 8, e67873. 10.1371/journal.pone.0067873.

18. Long, K.M., Whitmore, A.C., Ferris, M.T., Sempowski, G.D., McGee, C., Trollinger, B., Gunn, B., and Heise, M.T. (2013). Dendritic cell immunoreceptor regulates Chikungunya virus pathogenesis in mice. Journal of Virology 87, 5697–5706. 10.1128/jvi.01611-12.

19. Trimaglio, G., Sneperger, T., Raymond, B.B.A., Gilles, N., Näser, E., Locard-Paulet, M., Ijsselsteijn, M.E., Brouwer, T.P., Ecalard, R., Roelands, J., et al. (2024). The C-type lectin DCIR contributes to the immune response and pathogenesis of colorectal cancer. Sci. Rep. 14, 7199. 10.1038/s41598-024-57941-y.

20. Park, I., Goddard, M.E., Cole, J.E., Zanin, N., Lyytikäinen, L.-P., Lehtimäki, T., Andreakos, E., Feldmann, M., Udalova, I., Drozdov, I., et al. (2022). C-type lectin receptor CLEC4A2 promotes tissue adaptation of macrophages and protects against atherosclerosis. Nat Commun 13, 215. 10.1038/s41467-021-27862-9.

21. Weng, T.-Y., Li, C.-J., Li, C.-Y., Hung, Y.-H., Yen, M.-C., Chang, Y.-W., Chen, Y.-H., Chen, Y.-L., Hsu, H.-P., Chang, J.-Y., et al. (2017). Skin Delivery of Clec4a Small Hairpin RNA Elicited an Effective Antitumor Response by Enhancing CD8+ Immunity In Vivo. Mol Ther - Nucleic Acids 9, 419–427. 10.1016/j.omtn.2017.10.015.

22. Tokieda, S., Komori, M., Ishiguro, T., Iwakura, Y., Takahara, K., and Inaba, K. (2015). Dendritic cell immunoreceptor 1 alters neutrophil responses in the development of experimental colitis. BMC Immunology 16, 64. 10.1186/s12865-015-0129-5.

23. Meyer-Wentrup, F., Cambi, A., Joosten, B., Looman, M.W., Vries, I.J.M. de, Figdor, C.G., and Adema, G.J. (2009). DCIR is endocytosed into human dendritic cells and inhibits TLR8-mediated cytokine production. J Leukoc Biol 85, 518–525. 10.1189/jlb.0608352.

24. Zhao, X., Shen, Y., Hu, W., Chen, J., Wu, T., Sun, X., Yu, J., Wu, T., and Chen, W. (2015). DCIR negatively regulates CpG-ODN-induced IL-1β and IL-6 production. Molecular Immunology 68, 641–647. 10.1016/j.molimm.2015.10.007.

25. Nasu, J., Uto, T., Fukaya, T., Takagi, H., Fukui, T., Miyanaga, N., Nishikawa, Y., Yamasaki, S., Yamashita, Y., and Sato, K. (2020). Pivotal role of the carbohydrate recognition domain in self-interaction of CLEC4A to elicit the ITIM-mediated inhibitory function in murine conventional dendritic cells in vitro. Int Immunol 32, 673–682. 10.1093/intimm/dxaa034.

26. Meyer-Wentrup, F., Benitez-Ribas, D., Tacken, P.J., Punt, C.J.A., Figdor, C.G., Vries, I.J.M. de, and Adema, G.J. (2008). Targeting DCIR on human plasmacytoid dendritic cells results in antigen presentation and inhibits IFN-alpha production. Blood 111, 4245–4253. 10.1182/blood-2007-03-081398.

27. Lee, R.T., Hsu, T.-L., Huang, S.K., Hsieh, S.-L., Wong, C.-H., and Lee, Y.C. (2011). Survey of immune-related, mannose/fucose-binding C-type lectin receptors reveals widely divergent sugar-binding specificities. Glycobiology 21, 512–520. 10.1093/glycob/cwq193.

28. 28. Bloem, K., Vuist, I.M., Berk, M. van den, Klaver, E.J., Die, I.V., Knippels, L.M.J., Garssen, J., García-Vallejo, J.J., Vliet, S.J. van, and Kooyk, Y. van (2014). DCIR interacts with ligands from both endogenous and pathogenic origin. Immunology Letters 158, 33–41. 10.1016/j.imlet.2013.11.007.

29. 29. Bloem, K., Vuist, I.M., Plas, A.-J. van der, Knippels, L.M.J., Garssen, J., García-Vallejo, J.J., Vliet, S.J. van, and Kooyk, Y. van (2013). Ligand binding and signaling of dendritic cell immunoreceptor (DCIR) is modulated by the glycosylation of the carbohydrate recognition domain. Plos One 8, e66266. 10.1371/journal.pone.0066266.

30. Nagae, M., Ikeda, A., Hanashima, S., Kojima, T., Matsumoto, N., Yamamoto, K., and Yamaguchi, Y. (2016). Crystal structure of human dendritic cell inhibitory receptor (DCIR) C-type lectin domain reveals the binding mode with N-glycan. FEBS Lett. 10.1002/1873-3468.12162.

31. Massoud, A.H., Yona, M., Xue, D., Chouiali, F., Alturaihi, H., Ablona, A., Mourad, W., Piccirillo, C.A., and Mazer, B.D. (2014). Dendritic cell immunoreceptor: a novel receptor for intravenous immunoglobulin mediates induction of regulatory T cells. The Journal of allergy and clinical immunology 133, 853–63.e5. 10.1016/j.jaci.2013.09.029.

32. Yang, W., Ao, M., Hu, Y., Li, Q.K., and Zhang, H. (2018). Mapping the O-glycoproteome using site-specific extraction of O-linked glycopeptides (EXoO). Mol. Syst. Biol. 14, e8486. 10.15252/msb.20188486.

33. Gundry, R.L., Raginski, K., Tarasova, Y., Tchernyshyov, I., Bausch-Fluck, D., Elliott, S.T., Boheler, K.R., Eyk, J.E.V., and Wollscheid, B. (2009). The Mouse C2C12 Myoblast Cell Surface N-Linked Glycoproteome. Mol. Cell. Proteom. : MCP 8, 2555–2569. 10.1074/mcp.m900195-mcp200.

34. Khetarpal, S.A., Schjoldager, K.T., Christoffersen, C., Raghavan, A., Edmondson, A.C., Reutter, H.M., Ahmed, B., Ouazzani, R., Peloso, G.M., Vitali, C., et al. (2016). Loss of Function of GALNT2 Lowers High-Density Lipoproteins in Humans, Nonhuman Primates, and Rodents. Cell Metabolism 24, 234–245. 10.1016/j.cmet.2016.07.012.

35. Lageveen-Kammeijer, G.S.M., Haan, N. de, Mohaupt, P., Wagt, S., Filius, M., Nouta, J., Falck, D., and Wuhrer, M. (2019). Highly sensitive CE-ESI-MS analysis of N-glycans from complex biological samples. Nat. Commun. 10, 2137. 10.1038/s41467-019-09910-7.

36. Perusko, M., Grundström, J., Eldh, M., Hamsten, C., Apostolovic, D., and Hage, M. van (2024). The α-Gal epitope - the cause of a global allergic disease. Front. Immunol. 15, 1335911. 10.3389/fimmu.2024.1335911.

37. Galili, U. (2023). Antibody production and tolerance to the α-gal epitope as models for understanding and preventing the immune response to incompatible ABO carbohydrate antigens and for α-gal therapies. Front. Mol. Biosci. 10, 1209974. 10.3389/fmolb.2023.1209974.

38. Hatakeyama, T., Kamiya, T., Kusunoki, M., Nakamura-Tsuruta, S., Hirabayashi, J., Goda, S., and Unno, H. (2011). Galactose Recognition by a Tetrameric C-type Lectin, CEL-IV, Containing the EPN Carbohydrate Recognition Motif*. J. Biol. Chem. 286, 10305–10315. 10.1074/jbc.m110.200576.

39. Poget, S.F., Legge, G.B., Proctor, M.R., Butler, P.J.G., Bycroft, M., and Williams, R.L. (1999). The structure of a tunicate C-type lectin from polyandrocarpa misakiensis complexed with d-galactose11Edited by R. Huber. J. Mol. Biol. 290, 867–879. 10.1006/jmbi.1999.2910.

40. Bres, E.E., and Faissner, A. (2019). Low Density Receptor-Related Protein 1 Interactions With the Extracellular Matrix: More Than Meets the Eye. Front. Cell Dev. Biol. 7, 31. 10.3389/fcell.2019.00031.

41. Chen, J., Su, Y., Pi, S., Hu, B., and Mao, L. (2021). The Dual Role of Low-Density Lipoprotein Receptor-Related Protein 1 in Atherosclerosis. Front. Cardiovasc. Med. 8, 682389. 10.3389/fcvm.2021.682389.

42. Lillis, A.P., Duyn, L.B.V., Murphy-Ullrich, J.E., and Strickland, D.K. (2008). LDL Receptor-Related Protein 1: Unique Tissue-Specific Functions Revealed by Selective Gene Knockout Studies. Physiol. Rev. 88, 887–918. 10.1152/physrev.00033.2007.

43. Sizova, O., John, L.St., Ma, Q., and Molldrem, J.J. (2023). Multi-faceted role of LRP1 in the immune system. Front. Immunol. 14, 1166189. 10.3389/fimmu.2023.1166189.

44. Boucher, P., Gotthardt, M., Li, W.-P., Anderson, R.G.W., and Herz, J. (2003). LRP: Role in Vascular Wall Integrity and Protection from Atherosclerosis. Science 300, 329–332. 10.1126/science.1082095.

45. Bang, Y.-J., Hu, Z., Li, Y., Gattu, S., Ruhn, K.A., Raj, P., Herz, J., and Hooper, L.V. (2021). Serum amyloid A delivers retinol to intestinal myeloid cells to promote adaptive immunity. Science 373, eabf9232. 10.1126/science.abf9232.

46. Rauch, J.N., Luna, G., Guzman, E., Audouard, M., Challis, C., Sibih, Y.E., Leshuk, C., Hernandez, I., Wegmann, S., Hyman, B.T., et al. (2020). LRP1 is a master regulator of tau uptake and spread. Nature 580, 381–385. 10.1038/s41586-020-2156-5.

47. Ganaie, S.S., Schwarz, M.M., McMillen, C.M., Price, D.A., Feng, A.X., Albe, J.R., Wang, W., Miersch, S., Orvedahl, A., Cole, A.R., et al. (2021). Lrp1 is a host entry factor for Rift Valley fever virus. Cell 184, 5163–5178.e24. 10.1016/j.cell.2021.09.001.

48. Pencheva, N., Tran, H., Buss, C., Huh, D., Drobnjak, M., Busam, K., and Tavazoie, S.F. (2012). Convergent Multi-miRNA Targeting of ApoE Drives LRP1/LRP8-Dependent Melanoma Metastasis and Angiogenesis. Cell 151, 1068–1082. 10.1016/j.cell.2012.10.028.

49. Sedlacek, A.L., Younker, T.P., Zhou, Y.J., Borghesi, L., Shcheglova, T., Mandoiu, I.I., and Binder, R.J. (2019). CD91 on dendritic cells governs immunosurveillance of nascent, emerging tumors. JCI Insight 4, e127239. 10.1172/jci.insight.127239.

50. Mantuano, E., Brifault, C., Lam, M.S., Azmoon, P., Gilder, A.S., and Gonias, S.L. (2016). LDL receptor-related protein-1 regulates NFκB and microRNA-155 in macrophages to control the inflammatory response. P Natl Acad Sci Usa 113, 1369–1374. 10.1073/pnas.1515480113.

51. Luo, L., Wall, A.A., Tong, S.J., Hung, Y., Xiao, Z., Tarique, A.A., Sly, P.D., Fantino, E., Marzolo, M.-P., and Stow, J.L. (2018). TLR Crosstalk Activates LRP1 to Recruit Rab8a and PI3Kγ for Suppression of Inflammatory Responses. Cell Rep. 24, 3033–3044. 10.1016/j.celrep.2018.08.028.

52. Zurhove, K., Nakajima, C., Herz, J., Bock, H.H., and May, P. (2008). γ-Secretase Limits the Inflammatory Response Through the Processing of LRP1. Sci. Signal. 1, ra15. 10.1126/scisignal.1164263.

53. Pocivavsek, A., Mikhailenko, I., Strickland, D.K., and Rebeck, G.W. (2009). Microglial low-density lipoprotein receptor-related protein 1 modulates c-Jun N-terminal kinase activation. J. Neuroimmunol. 214, 25–32. 10.1016/j.jneuroim.2009.06.010.

54. Yang, L., Liu, C.-C., Zheng, H., Kanekiyo, T., Atagi, Y., Jia, L., Wang, D., N’songo, A., Can, D., Xu, H., et al. (2016). LRP1 modulates the microglial immune response via regulation of JNK and NF-κB signaling pathways. J. Neuroinflammation 13, 304. 10.1186/s12974-016-0772-7.

55. Gaultier, A., Arandjelovic, S., Niessen, S., Overton, C.D., Linton, M.F., Fazio, S., Campana, W.M., Cravatt, B.F., and Gonias, S.L. (2008). Regulation of tumor necrosis factor receptor-1 and the IKK-NF-κB pathway by LDL receptor–related protein explains the antiinflammatory activity of this receptor. Blood 111, 5316–5325. 10.1182/blood-2007-12-127613.

56. Xian, X., Ding, Y., Dieckmann, M., Zhou, L., Plattner, F., Liu, M., Parks, J.S., Hammer, R.E., Boucher, P., Tsai, S., et al. (2017). LRP1 integrates murine macrophage cholesterol homeostasis and inflammatory responses in atherosclerosis. Elife 6, e29292. 10.7554/elife.29292.

57. Basu, S., Binder, R.J., Ramalingam, T., and Srivastava, P.K. (2001). CD91 Is a Common Receptor for Heat Shock Proteins gp96, hsp90, hsp70, and Calreticulin. Immunity 14, 303–313. 10.1016/s1074-7613(01)00111-x.

58. Subramanian, M., Hayes, C.D., Thome, J.J., Thorp, E., Matsushima, G.K., Herz, J., Farber, D.L., Liu, K., Lakshmana, M., and Tabas, I. (2014). An AXL/LRP-1/RANBP9 complex mediates DC efferocytosis and antigen cross-presentation in vivo. J Clin Investigation 124, 1296–1308. 10.1172/jci72051.

59. Cui, H., Jiang, D., Banerjee, S., Xie, N., Kulkarni, T., Liu, R.-M., Duncan, S.R., and Liu, G. (2020). Monocyte-derived alveolar macrophage apolipoprotein E participates in pulmonary fibrosis resolution. Jci Insight 5. 10.1172/jci.insight.134539.

60. Mishra, A., Yao, X., Saxena, A., Gordon, E.M., Kaler, M., Cuento, R.A., Barochia, A.V., Dagur, P.K., McCoy, J.P., Keeran, K.J., et al. (2017). LRP-1 Attenuates House Dust Mite-induced Eosinophilic Airway Inflammation by Suppressing Dendritic Cell-mediated Adaptive Immune Responses. J Allergy Clin Immunol 142, 1066–1079.e6. 10.1016/j.jaci.2017.10.044.

61. Gardai, S.J., McPhillips, K.A., Frasch, S.C., Janssen, W.J., Starefeldt, A., Murphy-Ullrich, J.E., Bratton, D.L., Oldenborg, P.-A., Michalak, M., and Henson, P.M. (2005). Cell-Surface Calreticulin Initiates Clearance of Viable or Apoptotic Cells through trans-Activation of LRP on the Phagocyte. Cell 123, 321–334. 10.1016/j.cell.2005.08.032.

62. Gardai, S.J., Xiao, Y.-Q., Dickinson, M., Nick, J.A., Voelker, D.R., Greene, K.E., and Henson, P.M. (2003). By Binding SIRPα or Calreticulin/CD91, Lung Collectins Act as Dual Function Surveillance Molecules to Suppress or Enhance Inflammation. Cell 115, 13–23. 10.1016/s0092-8674(03)00758-x.

63. Vandivier, R.W., Ogden, C.A., Fadok, V.A., Hoffmann, P.R., Brown, K.K., Botto, M., Walport, M.J., Fisher, J.H., Henson, P.M., and Greene, K.E. (2002). Role of Surfactant Proteins A, D, and C1q in the Clearance of Apoptotic Cells In Vivo and In Vitro: Calreticulin and CD91 as a Common Collectin Receptor Complex. J Immunol 169, 3978–3986. 10.4049/jimmunol.169.7.3978.

64. Jégouzo, S.A.F., Feinberg, H., Dungarwalla, T., Drickamer, K., Weis, W.I., and Taylor, M.E. (2015). A Novel Mechanism for Binding of Galactose-terminated Glycans by the C-type Carbohydrate Recognition Domain in Blood Dendritic Cell Antigen 2*. J. Biol. Chem. 290, 16759–16771. 10.1074/jbc.m115.660613.

65. Karim, S., Leyva-Castillo, J.M., and Narasimhan, S. (2023). Tick salivary glycans – a sugar-coated tick bite. Trends Parasitol. 39, 1100–1113. 10.1016/j.pt.2023.09.012.

66. Chung, C.H., Mirakhur, B., Chan, E., Le, Q.-T., Berlin, J., Morse, M., Murphy, B.A., Satinover, S.M., Hosen, J., Mauro, D., et al. (2008). Cetuximab-induced anaphylaxis and IgE specific for galactose-alpha-1,3-galactose. N Engl J Med 358, 1109–1117. 10.1056/nejmoa074943.

67. Galili, U., Macher, B.A., Buehler, J., and Shohet, S.B. (1985). Human natural anti-alpha-galactosyl IgG. II. The specific recognition of alpha (1 3)-linked galactose residues. J. Exp. Med. 162, 573–582. 10.1084/jem.162.2.573.

68. Do, D.C., Yang, S., Yao, X., Hamilton, R.G., Schroeder, J.T., and Gao, P. (2017). N-glycan in cockroach allergen regulates human basophil function. Immun., Inflamm. Dis. 5, 386–399. 10.1002/iid3.145.

69. Lastrucci, C., Bénard, A., Balboa, L., Pingris, K., Souriant, S., Poincloux, R., Saati, T.A., Rasolofo, V., González-Montaner, P., Inwentarz, S., et al. (2015). Tuberculosis is associated with expansion of a motile, permissive and immunomodulatory CD16(+) monocyte population via the IL-10/STAT3 axis. Cell Research 25, 1333–1351. 10.1038/cr.2015.123.

70. Tzelepis, K., Koike-Yusa, H., Braekeleer, E.D., Li, Y., Metzakopian, E., Dovey, O.M., Mupo, A., Grinkevich, V., Li, M., Mazan, M., et al. (2016). A CRISPR Dropout Screen Identifies Genetic Vulnerabilities and Therapeutic Targets in Acute Myeloid Leukemia. Cell Rep. 17, 1193–1205. 10.1016/j.celrep.2016.09.079.

71. Accarias, S., Sanchez, T., Labrousse, A., Ben-Neji, M., Boyance, A., Poincloux, R., Maridonneau-Parini, I., and Cabec, V.L. (2019). Genetic engineering of Hoxb8-immortalized hematopoietic progenitors - a potent tool to study macrophage tissue migration. J. cell Sci. 133. 10.1242/jcs.236703.

72. Troegeler, A., Lastrucci, C., Duval, C., Tanne, A., Cougoule, C., Maridonneau-Parini, I., Neyrolles, O., and Lugo-Villarino, G. (2014). An efficient siRNA-mediated gene silencing in primary human monocytes, dendritic cells and macrophages. Immunology and Cell Biology 92, 699–708. 10.1038/icb.2014.39.

73. Bulteau, F., Thépaut, M., Henry, M., Hurbin, A., Vanwonterghem, L., Vivès, C., Roy, A.L., Ebel, C., Renaudet, O., Fieschi, F., et al. (2022). Targeting Tn-Antigen-Positive Human Tumors with a Recombinant Human Macrophage Galactose C-Type Lectin. Mol. Pharm. 19, 235–245. 10.1021/acs.molpharmaceut.1c00744.

74. Kuhn, B.T., Zimmermann, I., Egloff, P., Hürlimann, L.M., Hutter, C.A.J., Miscenic, C., Dawson, R.J.P., Seeger, M.A., and Geertsma, E.R. (2020). Biotinylation of Membrane Proteins for Binder Selections. Methods Mol. Biol. (Clifton, NJ) 2127, 151–165. 10.1007/978-1-0716-0373-4_11.

75. Burande, C.F., Heuzé, M.L., Lamsoul, I., Monsarrat, B., Uttenweiler-Joseph, S., and Lutz, P.G. (2009). A Label-free Quantitative Proteomics Strategy to Identify E3 Ubiquitin Ligase Substrates Targeted to Proteasome Degradation*. Mol. Cell. Proteom. 8, 1719–1727. 10.1074/mcp.m800410-mcp200.

76. Bouyssié, D., Hesse, A.-M., Mouton-Barbosa, E., Rompais, M., Macron, C., Carapito, C., Peredo, A.G. de, Couté, Y., Dupierris, V., Burel, A., et al. (2020). Proline: an efficient and user-friendly software suite for large-scale proteomics. Bioinformatics 36, 3148–3155. 10.1093/bioinformatics/btaa118.

77. Gaultier, A., Arandjelovic, S., Li, X., Janes, J., Dragojlovic, N., Zhou, G.P., Dolkas, J., Myers, R.R., Gonias, S.L., and Campana, W.M. (2008). A shed form of LDL receptor–related protein–1 regulates peripheral nerve injury and neuropathic pain in rodents. J. Clin. Investig. 118, 161–172. 10.1172/jci32371.

78. Marty, M.T., Baldwin, A.J., Marklund, E.G., Hochberg, G.K.A., Benesch, J.L.P., and Robinson, C.V. (2015). Bayesian Deconvolution of Mass and Ion Mobility Spectra: From Binary Interactions to Polydisperse Ensembles. Anal. Chem. 87, 4370–4376. 10.1021/acs.analchem.5b00140.

79. Kabsch, W. (2010). XDS. Acta Crystallogr. Sect. D 66, 125–132. 10.1107/s0907444909047337.

80. Liebschner, D., Afonine, P.V., Baker, M.L., Bunkóczi, G., Chen, V.B., Croll, T.I., Hintze, B., Hung, L.-W., Jain, S., McCoy, A.J., et al. (2019). Macromolecular structure determination using X-rays, neutrons and electrons: recent developments in Phenix. Acta Crystallogr. Sect. D, Struct. Biol. 75, 861–877. 10.1107/s2059798319011471.

81. Mirdita, M., Schütze, K., Moriwaki, Y., Heo, L., Ovchinnikov, S., and Steinegger, M. (2022). ColabFold: making protein folding accessible to all. Nat. Methods 19, 679–682. 10.1038/s41592-022-01488-1.

82. Emsley, P., Lohkamp, B., Scott, W.G., and Cowtan, K. (2010). Features and development of Coot. Acta Crystallogr. Sect. D: Biol. Crystallogr. 66, 486–501. 10.1107/s0907444910007493.

