## Supplemental information for "The human and mouse dendritic cell receptor DCIR binds to LRP1 N-glycans containing terminal galactoses, including the α-Gal antigen"

[1]Institut de Pharmacologie et de Biologie Structurale, IPBS, Université de Toulouse, CNRS, UPS, Toulouse, France
[2]Infrastructure Nationale de Protéomique, ProFI, FR 2048, Toulouse, France.
[3]Center for Proteomics and Metabolomics, Leiden University Medical Center, Leiden, The Netherlands
[4]Univ. Grenoble Alpes, CNRS, CEA, Institut de Biologie Structurale, Grenoble, France.
[5]Institut Universitaire de France, Paris, France.
[6]These authors contributed equally
[7]Present address: Biosciences Institute, Faculty of Medical Sciences, Newcastle University, Newcastle upon Tyne, United Kingdom.
[8]Present address: Centre for Inflammation Research, University of Edinburgh, Edinburgh BioQuarter, Edinburgh, United Kingdom.
[9]Present address: Institute for Clinical Chemistry and Laboratory Medicine, Faculty of Medicine, Technische Universität Dresden, Dresden, and National Center for Tumor Diseases (NCT), Partner Site Dresden, Dresden, German Cancer Research Center (DKFZ), Heidelberg, Germany.
[10]These senior authors contributed equally
[11]Lead contact

### SUPLEMENTARY TABLES

(in separate files)

Table S1: Proteins enriched by DCIR affinity pulldowns (related to Figures 1 and 2).

Table S2: Protein identifications from LRP1 immunoprecipitation (related to Figure 1).

Table S3: List and relative intensity of the N-glycans of human serum and mouse macrophage LRP1 $\alpha$-chains as analyzed by CE-ESI-MS (related to Figure 4).

Table S4: List of antibodies used for flow cytometry experiments.

### SUPLEMENTARY FIGURES

Figure S1: Flow cytometry gating strategy for the analysis of immune cells extracted from tissues of WT and mDCIR1-KO mice (related to Figure 1).

Figure S2: Flow cytometry analysis of mDCIR1+ cells in the lung, spleen and thymus of mice (related to Figure 1).

Figure S3: LRP1 is a main glycoprotein ligand of mDCIR1 (related to Figure 2)

Figure S4: Generation of LRP1-deficicent of mouse macrophages (related to Figures 2 and 3).

Figure S5: Transfection of siRNA abolishes hDCIR-ECD binding to mo-Mac along with LRP1 expression (related to Figure 3).

Figure S6: Glycosylation analysis of mouse and human LRP1 (related to Figure 4)

Figure S7: Structure comparison of hDCIR-ECD and hDCIR CTLD and sequence aligment of hDCIR with mouse DCIR homologues (related to Figures 5 and 6).

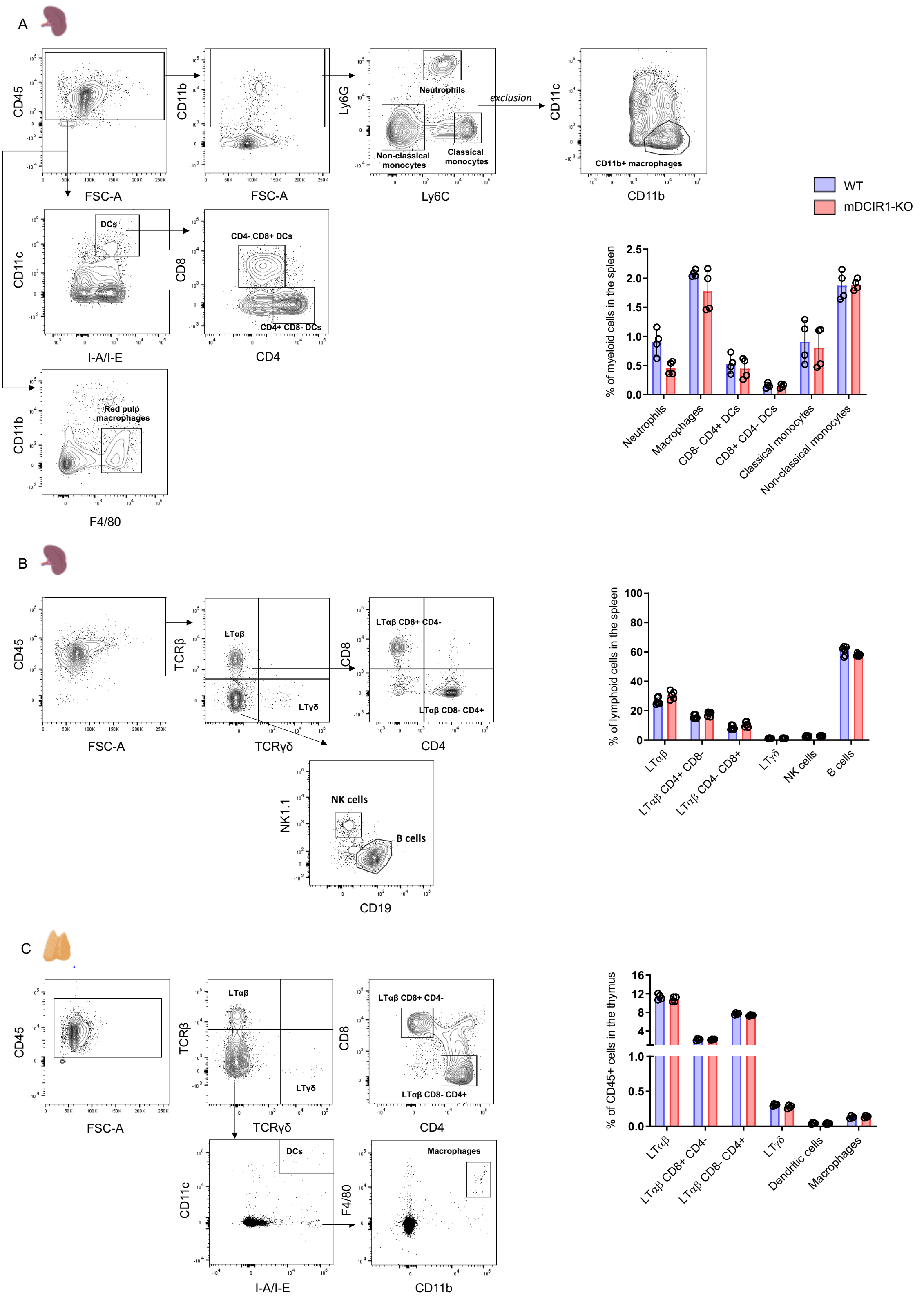
A
CD45
FSC-A
CD11b
Ly6G
Neutrophils
Non-classical monocytes
Classical monocytes
Ly6C
exclusion
CD11c
CD11b+ macrophages
CD11b
WT
mDCIR1-KO
DCs
I-A/I-E
CD8
CD4- CD8+ DCs
CD4+ CD8- DCs
CD4
Red pulp macrophages
F4/80
% of myeloid cells in the spleen
Neutrophils
Macrophages
CD8- CD4+ DCs
CD8+ CD4- DCs
Classical monocytes
Non-classical monocytes
B
TCRβ
LTαβ
LTγδ
TCRγδ
LTαβ CD8+ CD4-
LTαβ CD8- CD4+
NK1.1
NK cells
B cells
CD19
% of lymphoid cells in the spleen
LTαβ CD4+ CD8-
LTαβ CD4- CD8+
C
LTαβ CD8+ CD4-
LTαβ CD8- CD4+
Macrophages
% of CD45+ cells in the thymus
Dendritic cells

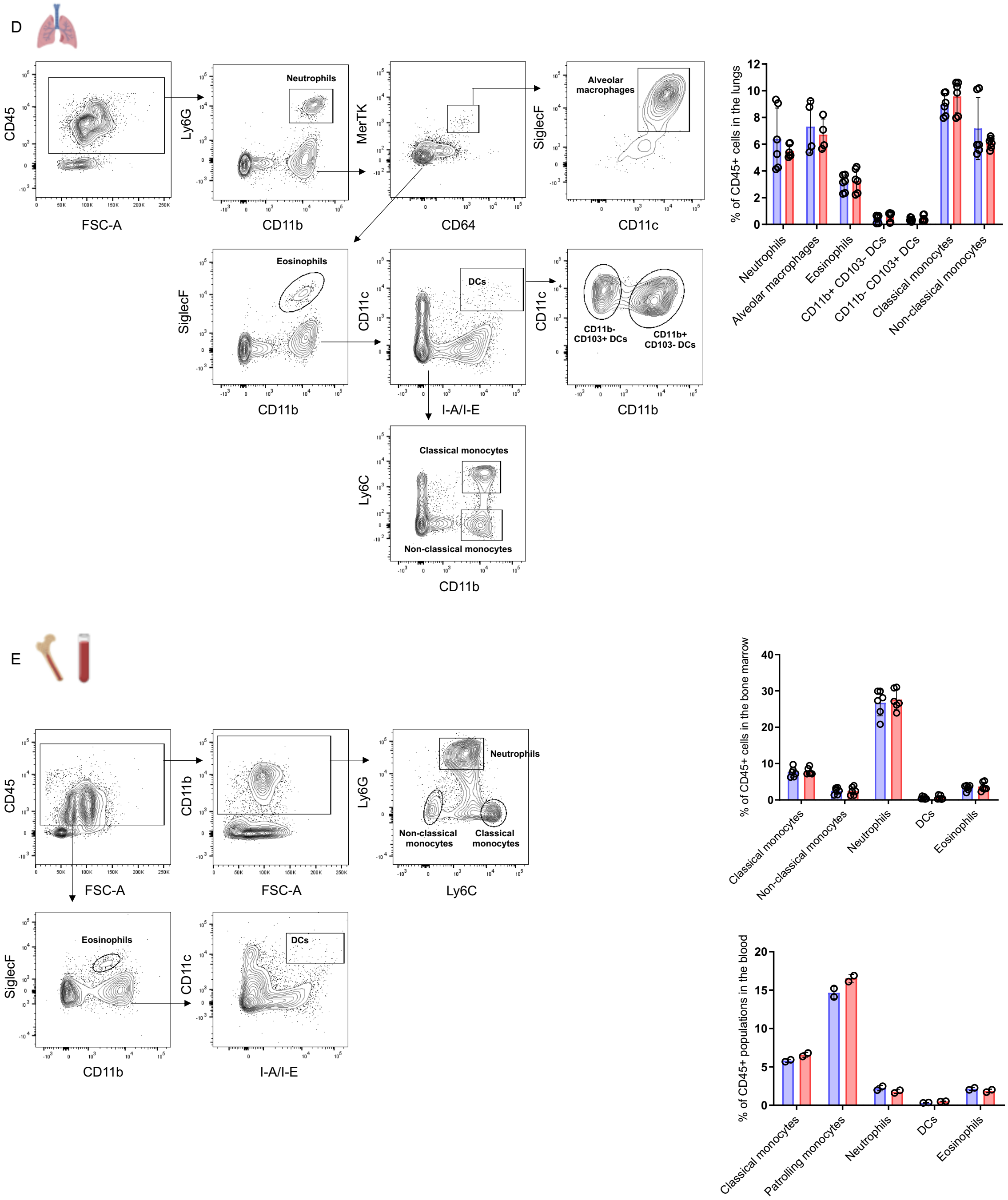


**Figure S1: Flow cytometry gating strategy for the analysis of immune cells extracted from tissues of WT and mDCIR1-KO mice (related to Figure 1).**

(A) Gating strategy (left panels) and percentages of myeloid cell subsets in the spleen of WT and mDCIR1-KO mice.

(B) Same as (A) but for lymphocytes.

(C) Gating strategy (left panels) and percentages of myeloid and lymphoid cell subsets in the thymus of WT and mDCIR1-KO mice.

(D) Same as (A) but for the lung.

(E) Same as (A) but for blood and bone marrow.

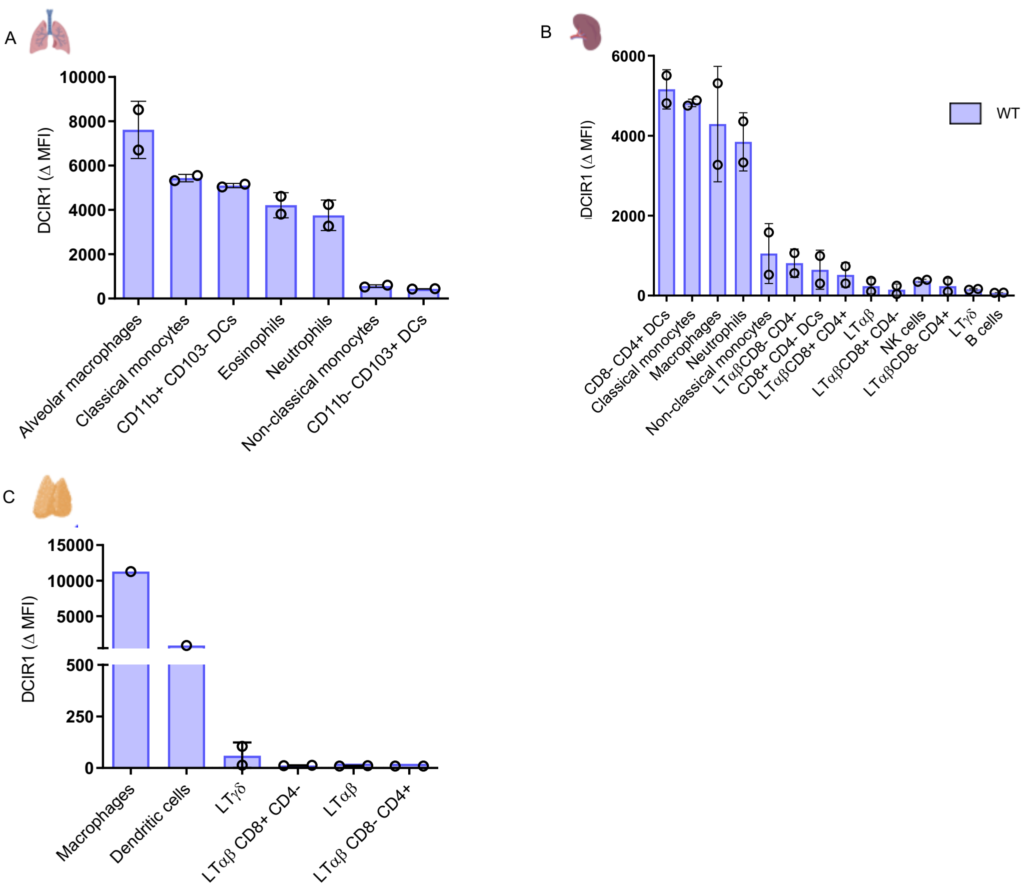


**Figure S2: Flow cytometry analysis of mDCIR1+ cells in the lung, spleen and thymus of mice (related to Figure 1).**

(A) ΔMFI (delta mean fluorescence intensity) of mDCIR1 staining in mouse lung.

(B) Same as (A) but for the spleen.

(C) Same as (A) but for the thymus.

**Figure S3: LRP1 is a main glycoprotein ligand of mDCIR1 (related to Figure 2)**

(A) Lectin blot examination of proteins extracted from BMDM and BMDC using biotinylated mDCIR1-ΔECD coupled to streptavidin-HRP.

(B) Schematic (Top) of the enrichment of mDCIR1 ligand(s) from BMDM by affinity purification using biotinylated mDCIR1-ECD (or in presence of EGTA chelating agent as negative control) coupled to streptavidin beads and their subsequent identification by proteomics (Created with BioRender.com). Volcano Plot (bottom) from quantitative mass spectrometry analyses of proteins differentially enriched (X-axis in log10) by affinity purification with mDCIR1-ECD versus mDCIR1-ECD with EGTA as a function of statistical significance (Y-axis in -log10). Dashed line marks the threshold limit (P-value = 0.5; Fold change = +/- 5).

(C) Western blot analysis of LRP1, LRPAP1 and THBS1 after affinity purification of BMDM proteins using either mDCIR1-ECD or mDCIR1-ΔECD.

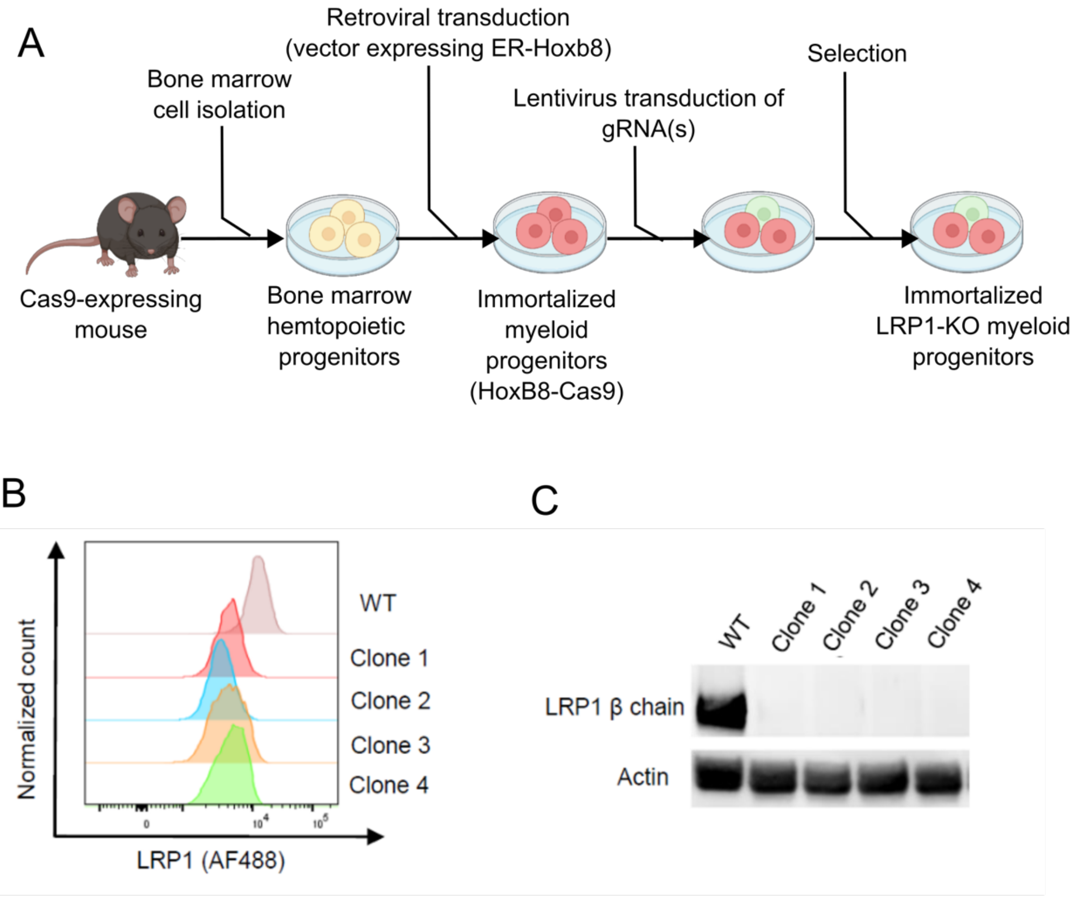


**Figure S4: Generation of LRP1-deficient mouse macrophages (related to Figures 2 and 3).**

(A) Schematic of the protocol used to generate LRP1-deficient HoxB8 myeloid progenitors. Created with BioRender.com

(B) Flow cytometry analysis of LRP1 β-chain in WT and four clones of LRP1-KO HoxB8-derived mouse macrophages

(C) Western blot analysis of LRP1 β-chain in WT and four clones of LRP1-KO HoxB8-derived mouse macrophages.

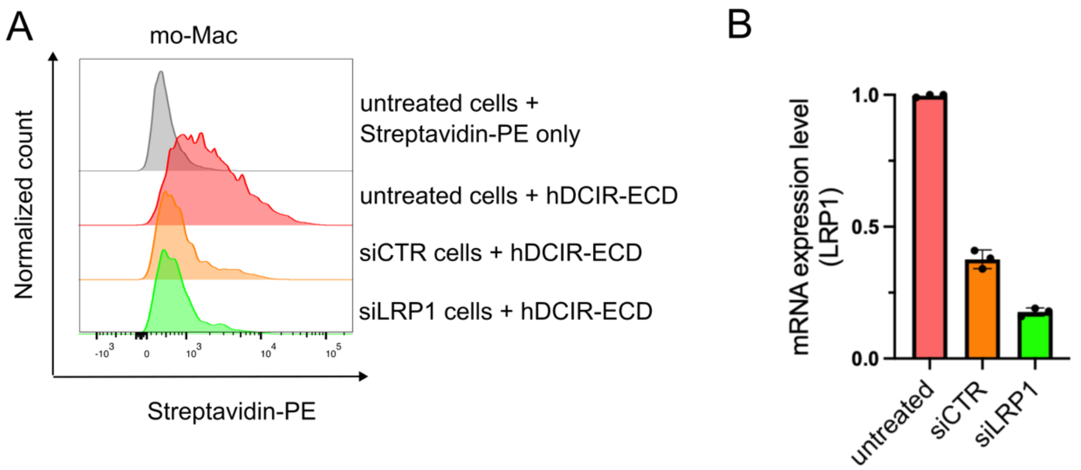


**Figure S5: Transfection of siRNA abolishes hDCIR-ECD binding to mo-Mac along with LRP1 expression (related to Figure 3).**

(A) Flow cytometry histograms of the binding of biotinylated hDCIR-ECD coupled to Streptavidin-PE to human blood monocyte-derived macrophages (mo-Mac) non-transfected, or transfected with either a non-targeted siRNA pool (siCTR) or siRNA against LRP1 (siLRP1). (B) mRNA expression level of LRP1 (in ddCT) normalized on GAPDH expression in mo-Mac) non-transfected, or transfected with either a non-targeted siRNA pool (siCTR) or siRNA against LRP1 (siLRP1).

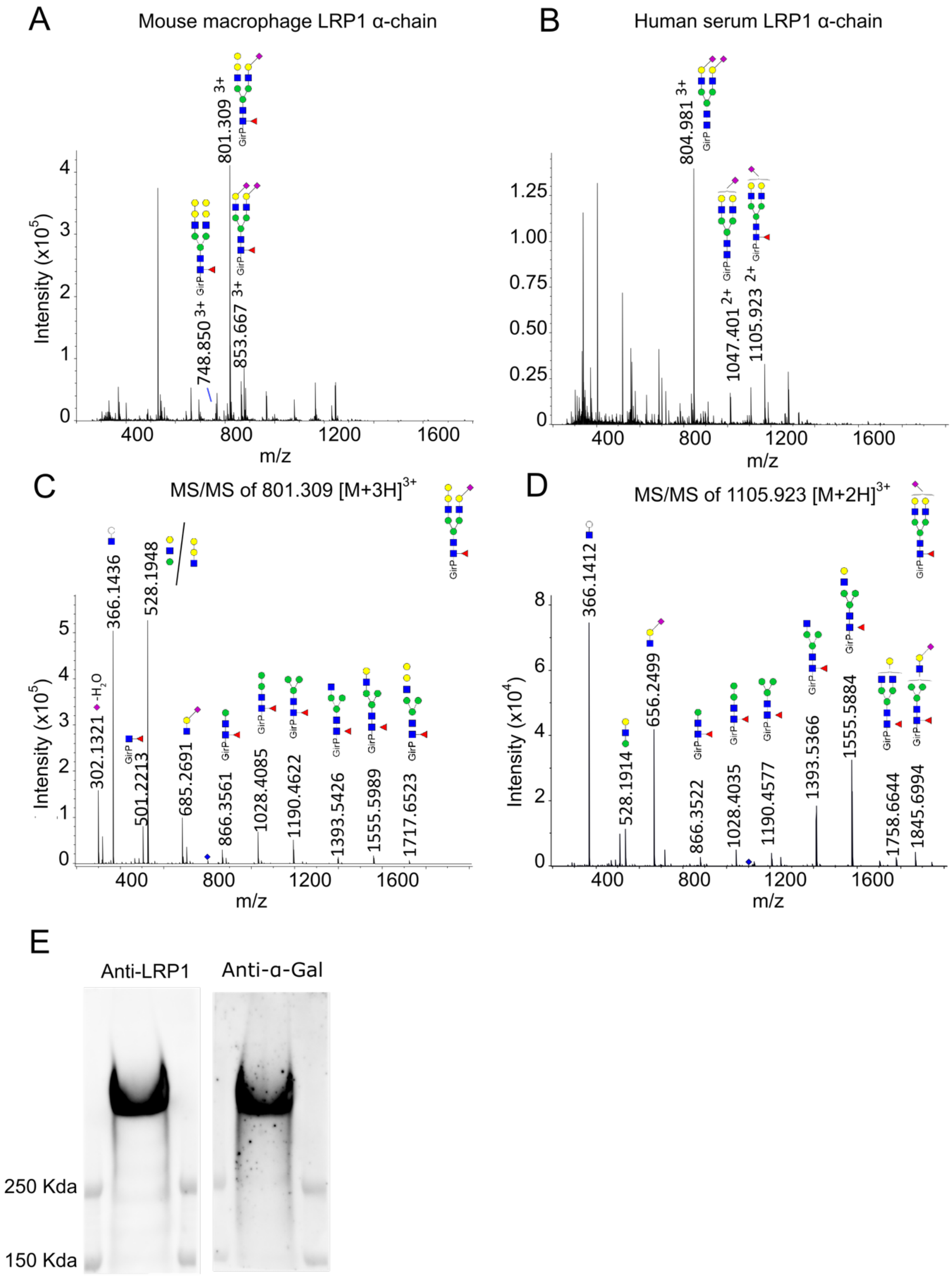
A
Mouse macrophage LRP1 α-chain
B
Human serum LRP1 α-chain
801.309 3+
748.850 3+
853.667 3+
Intensity (x10^5)
m/z
804.981 3+
1047.401 2+
1105.923 2+
C
MS/MS of 801.309 [M+3H]3+
D
MS/MS of 1105.923 [M+2H]3+
302.1321 -H2O
366.1436
528.1948
501.2213
685.2691
866.3561
1028.4085
1190.4622
1393.5426
1555.5989
1717.6523
366.1412
528.1914
656.2499
866.3522
1028.4035
1190.4577
1393.5366
1555.5884
1758.6644
1845.6994
Intensity (x10^4)
GirP
E
Anti-LRP1
Anti-α-Gal
250 Kda
150 Kda

**Figure S6: Glycosylation analysis of mouse and human LRP1 (related to Figure 4)**

(A) Summed mass spectrum (recorded between 40.1 and 43.6 min) of the CE-ESI-MS analysis of N-glycans from mouse macrophage LRP1 α-chain. N-glycans were analyzed after sialic acid derivatization (ethyl esterification and amidation) and reducing end labelling with Girard's reagent (GirP). Blue square: N-acetylglucosamine, green circle: mannose, yellow circle: galactose, red triangle: fucose, right pointing pink diamond: α2,6-linked N-acetylneuraminic acid/sialic acid, left pointing pink diamond: α2,3-linked N-acetylneuraminic acid/sialic acid.

(B) Summed mass spectrum (recorded between 40.3 and 43.3 min) of the CE-ESI-MS analysis of N-glycans from human serum LRP1 α-chain. N-glycans were derivatized and labelled as in described in (A).

(C). CE-ESI-MS/MS fragmentation spectra of the precursor ions at $801.309^{3+}$ of the derivatized and labelled N-glycans from mouse macrophage LRP1.

(D). CE-ESI-MS/MS fragmentation spectra of the precursor ions at $1105.923^{3+}$ of the derivatized and labelled N-glycans from human serum LRP1.

(E) Western blot confirmation of the presence of α-Gal epitope on mouse LPR1 α-chain.

A

B

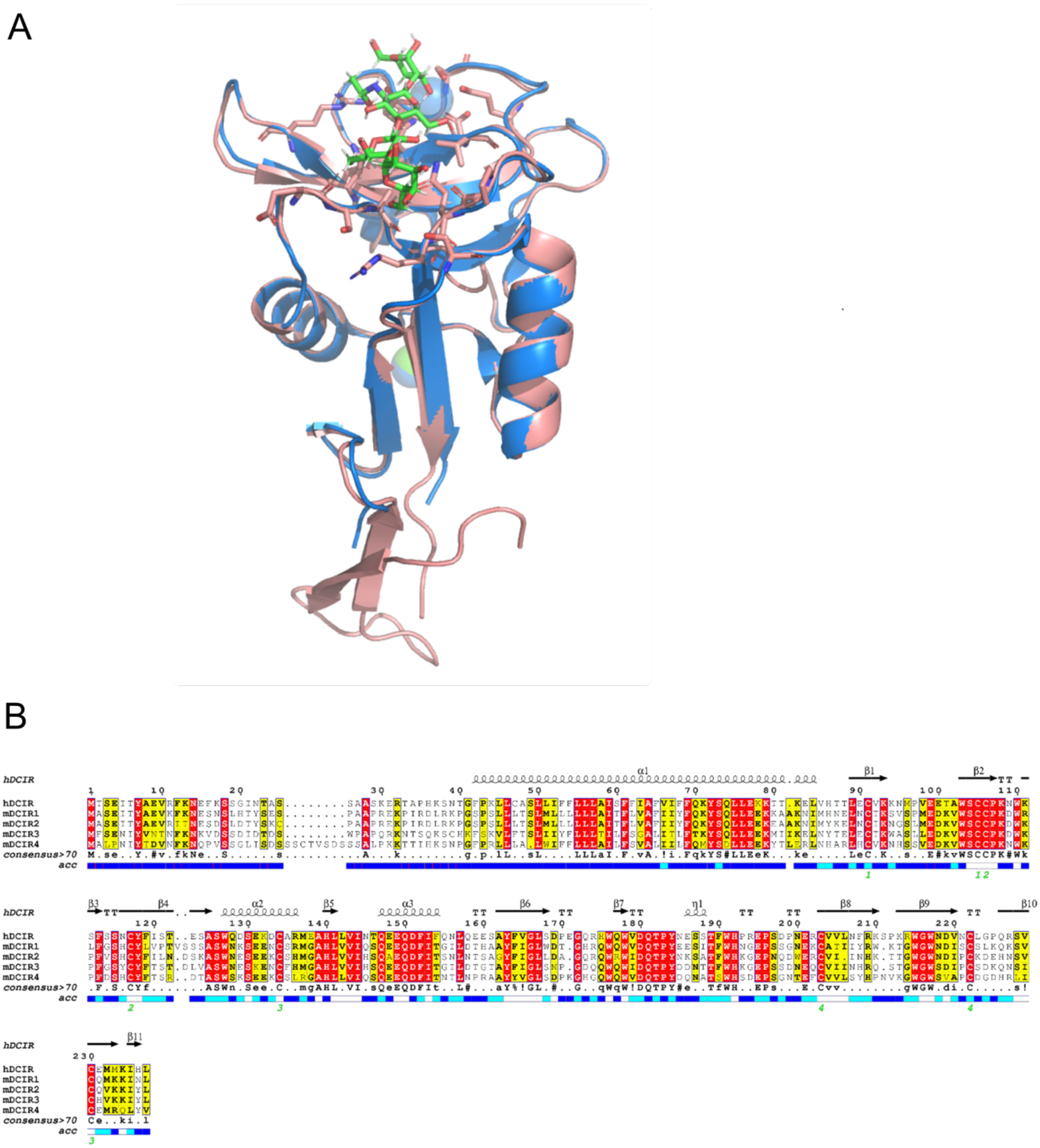


**Figure S7: Structure comparison of hDCIR-ECD and hDCIR-CTLD and sequence aligment of hDCIR with mouse DCIR homologues (related to Figures 5 and 6).**

(A) Structural superposition of hDCIR-ECD structure (in pink) and hDCIR-CTDL (*i.e.*, C-type lectin domain in blue; PDB 5b1x). Protein, carbohydrate, and calcium ions are shown in ribbon (pink/blue), stick (green/red), and sphere (light blue) models, respectively.

(B) Sequence alignment of human DCIR and mouse DCIR homologues (mDCIR1-4) as generated using ESPript 3.0 (https://espript.ibcp.fr/ESPript/ESPript/index.php). The secondary structure shown corresponds to that of human DCIR.
